# Fat cadherin cleavage releases a transcriptionally active nuclear fragment to regulate target gene expression

**DOI:** 10.1101/2025.05.08.652952

**Authors:** Jannette Rusch, Nattapon Thanintorn, Chikin Kuok, Yonit Tsatskis, Huayun Hou, Michael Wilson, Cole Julick, Helen McNeill

## Abstract

The conserved atypical cadherin *fat* (*ft*) controls cellular processes such as growth, planar cell polarity, and mitochondrial function, in organisms ranging from fruit flies to mammals. Working at the apical-junctional plasma membrane the intracellular domain of the Ft protein, FtICD, binds to and regulates components of the Hippo and PCP pathways. Unexpectedly, we show that FtICD is present in the nucleus in cultured cells as well as in embryonic and larval tissues, and identify nuclear localization and nuclear export signals in FtICD required for this localization. We show that membrane-bound FtICD is cleaved and enters nuclei *in vivo*. Using endogenously tagged Ft as well as overexpressed FtICD we conducted ChIP-seq experiments and identified putative Ft targets including genes involved in signaling pathways, chromatin organization, pattern formation, and neural development. RNAseq demonstrates that some of these genes are differentially regulated in *ft* mutants. We observe strong correlations of Ft binding regions with peaks for other factors such as DREF and BEAF-32, as well as the Hpo pathway components Yorkie (Yki) and Scalloped (Sd), suggesting that Ft may act in conjunction with these factors to regulate gene expression. Supporting this hypothesis, we found that Ft can physically interact with both Yki and Sd in co-immunoprecipitation experiments in S2 cells. We propose that the modulation of Hippo pathway activity constitutes one of the nuclear functions of Ft, complementing its established function as an upstream regulator of Hippo signaling.

## Introduction

Control of tissue growth and homeostasis is a crucial process that all multicellular organisms must accomplish in order to develop and function correctly. A number of pathways that contribute to this control have been identified, among them the conserved Hippo pathway (Edgar, 2006; Kango-Singh & Singh, 2009; Reddy & Irvine, 2008; Zheng & Pan, 2019). Its core components are two kinases, Hippo (Hpo) and Warts (Wts), which control the nuclear localization and activity of a transcriptional co-activator, Yorkie (Yki) (Dong et al., 2007; Huang et al., 2005; Oh & Irvine, 2008; Ren et al., 2010). Yki, together with the DNA-binding transcription factor Scalloped (Sd), regulates the transcription of Hpo target genes including genes that control proliferation and apoptosis, such as *diap1* and *cycE* (S. Wu et al., 2008; L. Zhang et al., 2008). The atypical cadherin family member Fat (Ft) has emerged as an upstream regulator of this pathway (Bennett & Harvey, 2006; Cho et al., 2006; Pan, 2010; Silva et al., 2006; Willecke et al., 2006). Ft is a conserved giant cell adhesion molecule with a single transmembrane domain and an intracellular domain containing multiple functional and conserved domains. Ft is localized to apical junctions and regulates the Hpo pathway through interactions with pathway members such as Expanded (Ex) (Fulford et al., 2023; Silva et al., 2006) and Dachs (D) (Mao et al., 2006), and interacts via cadherin-cadherin binding with another atypical cadherin, Dachsous (Ds) (Blair & McNeill, 2018; Fulford & McNeill, 2020; Irvine & Harvey, 2015). In addition to its role in the Hpo pathway, Fat is also part of an essential pathway regulating a form of tissue organization, planar cell polarity (PCP), (Matis & Axelrod, 2013; Strutt & Strutt, 2021) which, coordinately with tissue growth, ensures the proper formation and patterning of structures such as wings and eyes. Finally, a fragment of Fat is imported into mitochondria where it binds directly to a core component of complex I (CI) of the electron transport chain, Ndufv2, stabilizing the CI holoenzyme as well as complex V (CV), and promoting oxidative phosphorylation crucial to proper development (Sing et al., 2014).

A number of proteolytic cleavage events of the Fat protein have been demonstrated and are necessary for Fat to fulfill its functions. These include the cleavage of the 560 kD full-length protein into 450 kD and 110 kD fragments, which form a stable complex at the membrane (Feng & Irvine, 2009a; Sopko et al., 2009), as well as an intracellular cleavage generating a ∼68 kD fragment imported into mitochondria, Ft^mito^ (Sing et al., 2014). The nature of the proteases carrying out these cleavages are currently unknown. However, in the case of Ft^mito^, it has been shown that the cleavage only takes place if the Ft precursor is membrane-anchored (Sing et al., 2014), suggesting that the responsible proteases are also membrane associated.

In a previous study aimed at identifying potential interactors of Ft, the transcription corepressor Atrophin (atro) was found to interact physically and genetically with Ft (Fanto et al., 2003), raising the possibility that a fragment of Ft could be imported into the nucleus. The suggestion of a nuclear function would add a fourth mechanism of action for Ft, in addition to its previously described roles in regulating growth *via* the Hpo pathway, regulating PCP, and promoting mitochondrial function. Here, we describe our findings that an intracellular fragment of Ft can be transported into the nucleus both in cultured cells and in tissues, and that this nuclear fragment can bind DNA at specific sites and regulate transcription. A subset of these sites are direct Yki/Sd targets, and we show that they form a complex. This novel function for Ft extends its roles in regulation of cell growth and tissue organization.

## Results

### Overexpressed Ft intracellular domain is present in the nucleus

To examine the possibility of nuclear localization of Ft, we examined the expression of over-expressed, C-terminally HA-tagged Ft constructs. The Ft intracellular domain (ftICD, see Fig. 1H) was expressed in S2 cells and visualized by co-immunostaining with HA and the nuclear envelope marker lamin. Ft-ICD-HA was significantly enriched in the nucleus (Fig. 1 A-C). To extend this finding, we examined tagged Ft constructs (Fig. 1H) *in vivo*. Full-length Ft, extracellular domain-deleted Ft (ftμECD), and Ft intracellular domain (FtICD) constructs were expressed in eye imaginal discs under GMR-Gal4 control, and wandering L3 larvae immunostained for HA. GMR-Gal4 drives expression in cells posterior to the morphogenetic furrow (Ellis et al., 1993). As expected, full-length Ft is localized at the apical membrane (Fig. 1D). When most of the extracellular domain is removed but the transmembrane and intracellular domains are retained (FtμECD), the protein remains at the cell membrane but is not restricted to the apical membrane (Fig. 1E). In contrast, FtICD is cytoplasmic in cells near the morphogenetic furrow, when it is first expressed, and then becomes localized to the nucleus as cells leave the furrow and mature. (Fig. 1F, schematic of eye disc morphology shown in Fig. 1G). Co-staining with the DNA marker propidium iodide (PI) (Suzuki et al., 1997) further shows that FtICD-HA colocalizes with DNA (Fig. 1I-K), while co-staining with the nucleolus marker fibrillarin indicates that FtICD is excluded from the nucleolus (Fig. S1A-B).

**Fig. 1:**
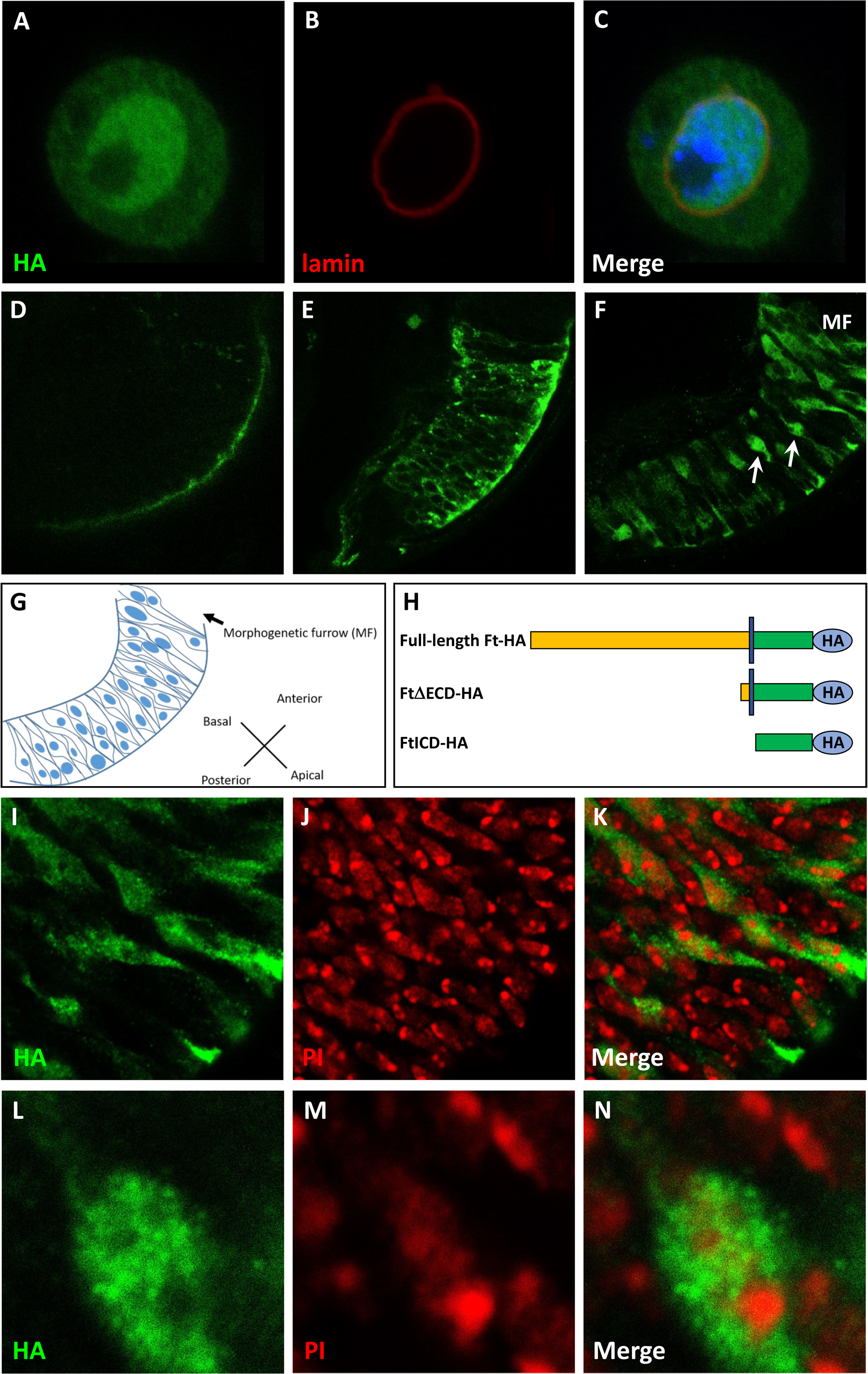
The intracellular domain of ft (ftICD) can localize to the nucleus. (A-C), S2 cell transfected with ftICD-HA (green). The nucleus is labeled with DAPI (blue) and outlined with lamin (red). ftICD can be seen accumulating in the nucleus. HA-tagged full-length ft (D), ftμECD (E) and ftICD (F) were overexpressed in 3^rd^ instar larval eye discs using GMR-Gal4 and stained for HA. Full-length ft is restricted to the apical surface of the tissue (D) while ftμECD is seen throughout the cell membrane (E). FtICD can be seen localizing to nuclei (arrows) in areas posterior to the morphogenetic furrow (MF) (F). (G), schematic outlining the morphology of the eye disc in (F). (H), diagram of constructs used in (D-F). (I-N), co-staining of GMR>FtICD expressing eye discs with HA to label ftICD (green, I and L) and propidium iodide (red, J and M) to label DNA. FtICD co-localizes with DNA but is excluded from nucleoli as seen in the closeup (N).

Significantly, endogenous Ft protein can be detected localizing to DNA in polytene chromosomes in *wild-type* (Fig. S1 C-F), but not *ft* mutant (Fig. S1 G-I), salivary gland nuclei. Some of the Ft-positive polytene bands (Fig. S1 D and F) overlap with RNA polII bands (Fig. S1 C and F), suggesting that Ft may play a role in transcription.

Endogenous Ft is not detectable by immunostaining in nuclei in any tissue that we examined, indicating that the levels of nuclear Ft must be very low. However, this does not preclude a role for nuclear Ft, as this scenario is reminiscent of Notch where undetectable levels of Notch in the nucleus are nevertheless active (Crittenden et al., 1994; Fehon et al., 1991; Fortini et al., 1993; Nye et al., 1994; Roehl & Kimble, 1993; Schroeter et al., 1998).

### Proteolytic cleavage and nuclear import of Ft protein

Translocation of an intracellular fragment of Ft to the nucleus requires a proteolytic cleavage event to release it from its membrane-bound full-length precursor. There is precedence for Ft cleavage as it has previously been shown that full-length Ft is subject to a constitutive proteolytic cleavage which generates a heterodimer of N and C-terminal fragments of Ft (Feng & Irvine, 2009b; Sopko et al., 2009), as well as an intracellular cleavage that generates a 68kD fragment which enters mitochondria (Sing et al., 2014). To detect such a cleavage event, a fly strain carrying a membrane-bound Ft protein (FtμECD) fused to a DNA binding and transactivation domain (Gal4-VP16) under control of the tubulin promoter, Tub-FtμECD-Gal4-VP16 (Fig. 2E), was crossed to a reporter strain (UAS-lacZ) and stained for βgal. In both wing and eye discs robust βgal activity is detected when both transgenes are present (Fig. 2B and 2D), while no signal is seen with the reporter construct only (Fig. 2A and 2C). Since the Gal4-VP16 domain-containing protein is tethered to the cell membrane and therefore unable to bind DNA and activate transcription, this result strongly suggests that a proteolytic cleavage event must have occurred to release an intracellular fragment to enter the nucleus. It is currently unknown which protease executes this cleavage.

**Fig. 2:**
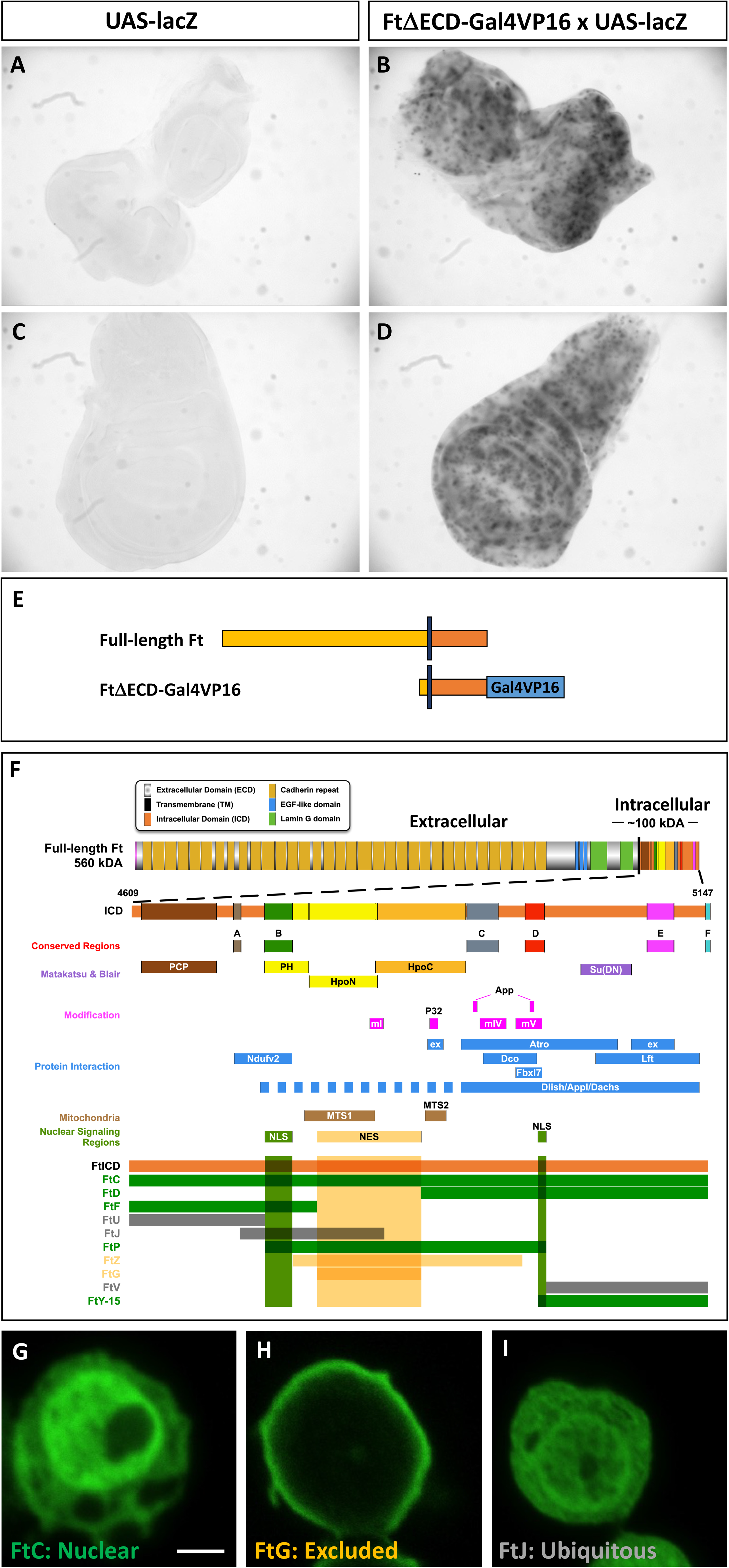
The ft intracellular domain can be cleaved *in vivo* and contains sequences required for nuclear import and export. β-gal stainings of eye (A, B) and wing (C, D) discs of animals carrying the indicated transgenes. Membrane-bound FtμECD-Gal4VP16 is cleaved to enter the nucleus and activate the UAS-lacZ reporter (B, D). (E), Diagram of the construct used in (B) and (D). (F), Schematic of the Ft protein with focus on the intracellular region (orange bar) indicating conserved regions (red) and functional domains as identified by Matakatsu and Blair (2012) (purple). The constructs represented by the bars (with HA tags) were transiently transfected into S2 cells and stained with antibodies to HA. Signal distribution was classified as nuclear-enriched (green bars), nuclear-excluded (yellow bars), or ubiquitous (gray bars). Representative examples for the different classes are shown in (G) – (I). Scale bar, 2 μm.

The molecular weight of the Ft intracellular domain (estimated at 58kD) (Stothard, 2018) is too large to allow passive diffusion into the nucleus (Mohr et al., 2009). We examined FtICD for the presence of nuclear localization (NLS) or nuclear export (NES) signals using computational tools (see Methods), however no predicted NLS or NES were found. To identify regions within FtICD that are necessary and sufficient for nuclear import and export, a series of truncations was made, transiently transfected into S2 cells, and analyzed by immunofluorescence (Fig. 2F-I and Fig. S2). Constructs were classified as nuclear-enriched (Fig. 2G), nuclear excluded (Fig. 2H) or ubiquitous (Fig. 2I). This led to the identification of 2 regions promoting nuclear localization (NLS) and one region promoting nuclear export (NES) (Fig. 2F). Notably, these sequences overlap with previously reported functional and conserved regions (Matakatsu & Blair, 2012): NLS1 overlaps with PH/Conserved Region B, NLS2 overlaps with Conserved Region D, and NES overlaps with HpoN and HpoC.

### FtICD can activate the hippo pathway target *diap1*

Previous studies have shown that Ft regulates target genes of the Hippo pathway such as *cyclinE* and *death-associated inhibitor of apoptosis 1* (*diap1*) (Silva et al., 2006; Willecke et al., 2006). To investigate whether Ft may be directly involved in this transcriptional regulation we made use of a short element from the *diap1* enhancer region that was previously identified as a Hippo Response Element (HRE, see Fig. 3E) (S. Wu et al., 2008). The diap1 HRE is a 32 bp fragment that activates transcription of a lacZ reporter in the absence of *hpo* signaling, such as in *hpo* mutant clones in imaginal discs (S. Wu et al., 2008). This element contains a binding site for the TEAD transcription factor Scalloped (Sd), which is necessary for its activation, as an HRE with a mutated Sd site (HREm6) is not activated in *hpo* clones (S. Wu et al., 2008). We examined HRE and HREm6 in *hpo* mutant MARCM clones (positively marked by GFP) with and without overexpressed FtICD in L3 eye and wing discs (Fig.3 and Fig. S3). As expected, HRE is activated in *hpo* clones both with and without overexpressed FtICD in wing and eye discs (Fig. 3A-D” and Fig. S3A-D”). When HREm6 was used there was no detectable expression of βgal in *hpo* clones in either wing (Fig. 3C-C”) or eye (Fig. S3C-C”) discs. However, in the presence of FtICD, HREm6 could be activated in some wing disc (Fig. 3D-D”) but not eye disc clones (Fig. S3D-D”). While these results do not prove that Ft acts on HRE directly, they support the notion that Ft can function in the nucleus.

**Fig. 3:**
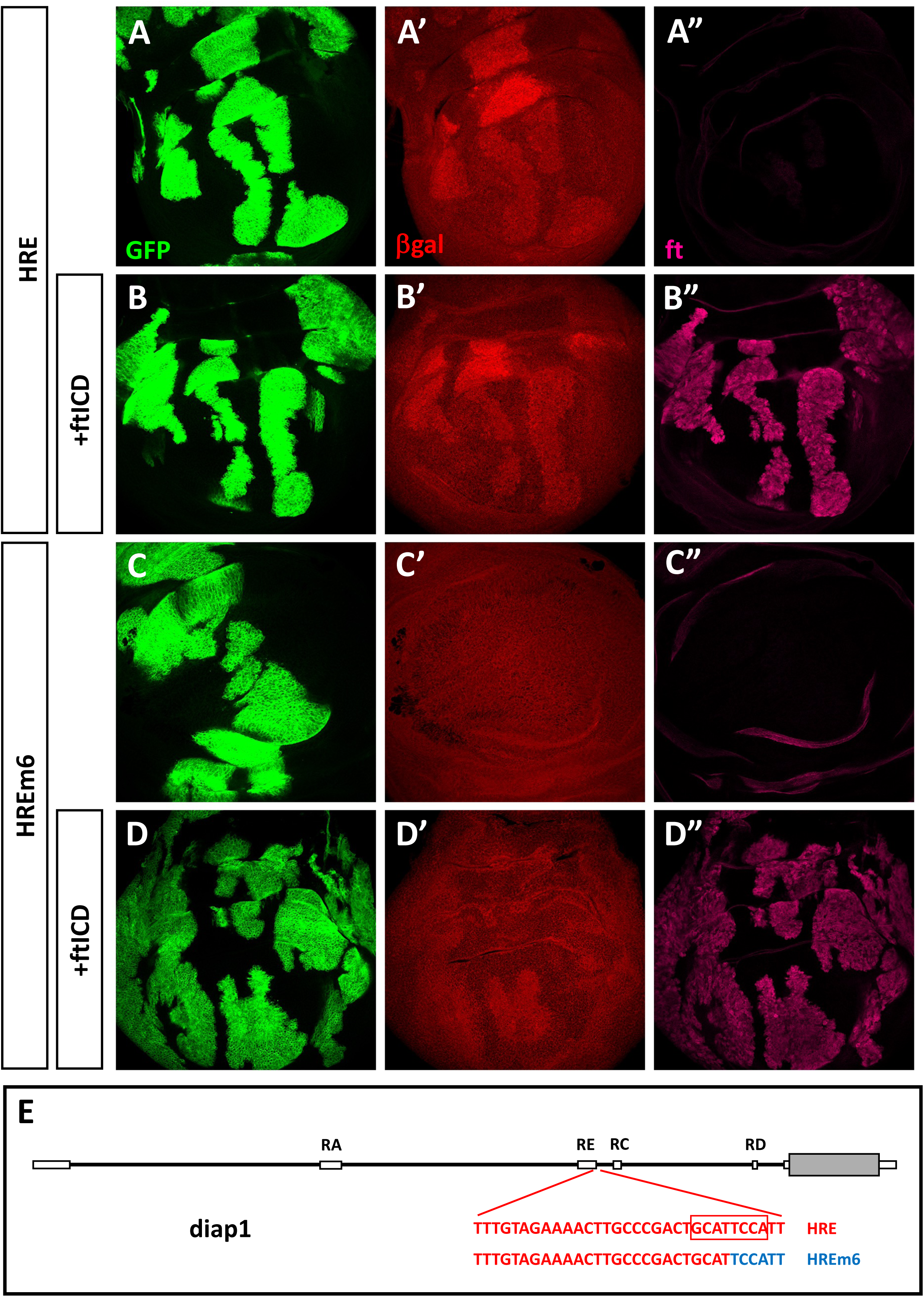
ftICD can activate the hippo pathway target *diap1*. *hpo^42-47^* mutant clones (marked by GFP expression in green) were generated in wing discs in larvae carrying either the HRE (A, B) or HREm6 (C, D) reporter. Larvae also carrying the UAS-ftICD transgene are shown in (B) and (D). Discs were stained with antibodies to βgal (red) and ft (magenta). Wild-type HRE is activated in *hpo* clones (A, B), whereas HREm6, which lacks the sd binding site, is not activated (C). When ftICD is overexpressed in the *hpo* clones, HREm6 can be activated in some wing discs (D). (E) Schematic of the diap1 locus indicating the location of HRE in the diap1 enhancer region. The Sd binding site is indicated by the box, and the bases mutated in HREm6 are indicated in blue.

### FtICD functions independently of *wts* and *yki*

The known functions of Ft that are mediated by its interactions with the Hippo pathway are dependent on its localization at the cell membrane, where it regulates pathway components such as Dachs and Expanded (Fulford et al., 2023; Matakatsu & Fehon, 2024). Consistent with this notion, overexpression of FtICD, which is not membrane-tethered is unable to rescue *ft* mutant phenotypes (Matakatsu & Blair, 2012). Mutations in Hippo pathway components such as *wts* and *yki* have strong effects on tissue growth, and while both are lethal as homozygotes, small changes in adult wing size can be observed in heterozygous animals (Wada et al., 2021). We aimed to determine whether FtICD could act independently of *wts* and/or *yki*, indicating a function separate from its role in regulating the Hippo pathway. Overexpression of FtICD causes slight wing undergrowth and a reduction in the crossvein distance, while *wts* heterozygotes display slightly larger wings (Fig. 4A-D). When FtICD is overexpressed in the posterior compartment of the developing wing disc in the context of a *wts* heterozygous background, wing size and crossvein distance resemble those of animals overexpressing FtICD in a *wild-type* background (Fig. 4A-D, wing parameters defined in Fig. S4). Thus, the nuclear form of Ft, Ft-ICD, is epistatic to wts. *yki* heterozygous wings are slightly smaller than *wild-type*, but their crossvein distance is not significantly different (Fig. 4A-D). When FtICD is overexpressed in the posterior compartment of *yki* heterozygous wings, their size is decreased further (Fig. 4A-C), and their crossvein distance is reduced to values similar to those seen in animals overexpressing FtICD alone (Fig. 4A, D). Analysis of the degree of roundness revealed that *wts* heterozygous wings are notably round while overexpression of FtICD did not cause increased roundness (Fig. 4E). Remarkably, overexpression of FtICD in a *wts* heterozygous background significantly increased roundness beyond both FtICD or *wts* het alone. Together, these results suggest that FtICD can function to affect tissue growth *via* a mechanism that does not involve the Hippo signaling cascade *per se* and supports a possible nuclear role for Ft in addition to or in parallel with Yki to regulate PCP. Sequestration of Yki could explain the effects of FtICD overexpression on wing size, so we also examined the effects on planar cell polarity in the fly eye (Fig. 4F and 4G). Remarkably, expression of FtICD in the eye leads to significant alterations in PCP, with dorsal-ventral inversions in polarity, similar to those seen in *ft* and *ds* mutant tissue, suggesting nuclear FtICD has functions in PCP organization.

**Fig. 4:**
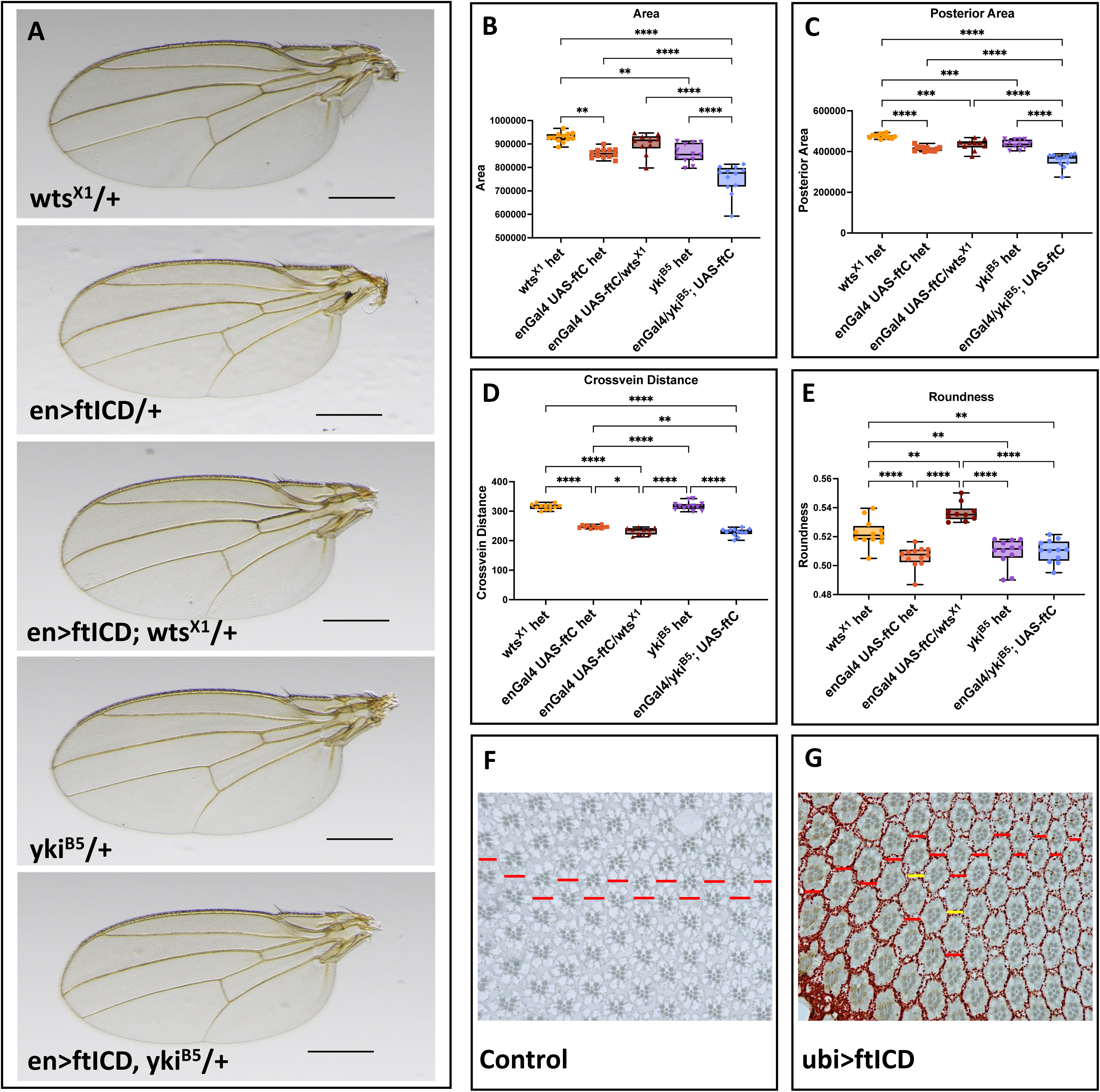
ftICD is epistatic to *wts* and *yki*. Wings from adult female flies of the indicated genotypes were measured and analyzed. (A) Representative images of adult wings. (B) Total wing area measurements. (C) Area of the posterior wing, defined as area below L4. (D) Distance between the anterior and posterior crossveins. (E) Wing roundness, defined as (4 * area)/ (ν * major axis length^2^). Scale bar, 500 μm. (F) and (G) Tangential sections of control (F) and FtICD overexpressing (G) eyes. The red horizontal lines indicate where dorsal and ventral forms meet forming the equator. In animals overexpressing ftICD there are ectopic mini-equators represented by the red lines, and yellow lines showing where the ommatidial inversions result in inappropriate “inverted” equators.

### Potential targets of overexpressed FtICD in larval imaginal tissue

Since overexpressed FtICD is readily detected in nuclei, and since Ft’s function upstream of the Hippo pathway depends on its membrane-proximal location (Matakatsu & Blair, 2006), we expressed FtICD *in vivo* to identify potential targets of Ft. We performed ChIP-seq on L3 eye-brain complexes expressing C-terminally FLAG-tagged FtICD (ftC-3xFLAG) driven by ubi-Gal4. In these animals, high levels of expression are seen across larval tissues, shown for eye discs and brains (Fig. S5). As there are no known DNA binding motifs in FtICD, we reasoned that Ft may interact with DNA indirectly. We used a 2-step crosslinking protocol with an EGS step preceding the standard formaldehyde crosslinking step as with chromatin associated proteins and used antibodies to the FLAG tag. ChIP experiments were done in triplicate. Peaks were selected based on their fold enrichment (FE) over controls, and a cutoff of FE>2 was used. We found that a large fraction of reproducible peaks are located at promoter regions, mostly within 1kb of the transcription start site (Fig. 5A). GO term analysis focusing on signaling revealed enrichment of several pathways that Ft plays central roles in (planar polarity, Hippo signaling, Fig. 5B). 2 examples of putative Ft targets that are involved in the Hippo pathway, *dachs* (*d*) and *discs overgrown* (*dco*) are shown in Fig. 5C and 5D. The Hippo target gene, *diap1*, was also identified as a putative Ft target (Fig. S5). Interestingly, we found that one of the *diap1* Ft ChIP peaks overlaps the HRE sequence (orange box in Fig. S5E) which is located in an intron/5’ of the diap1-RC isoform (see Fig. 3E), supporting the idea that Ft may bind to this regulatory element.

**Fig. 5:**
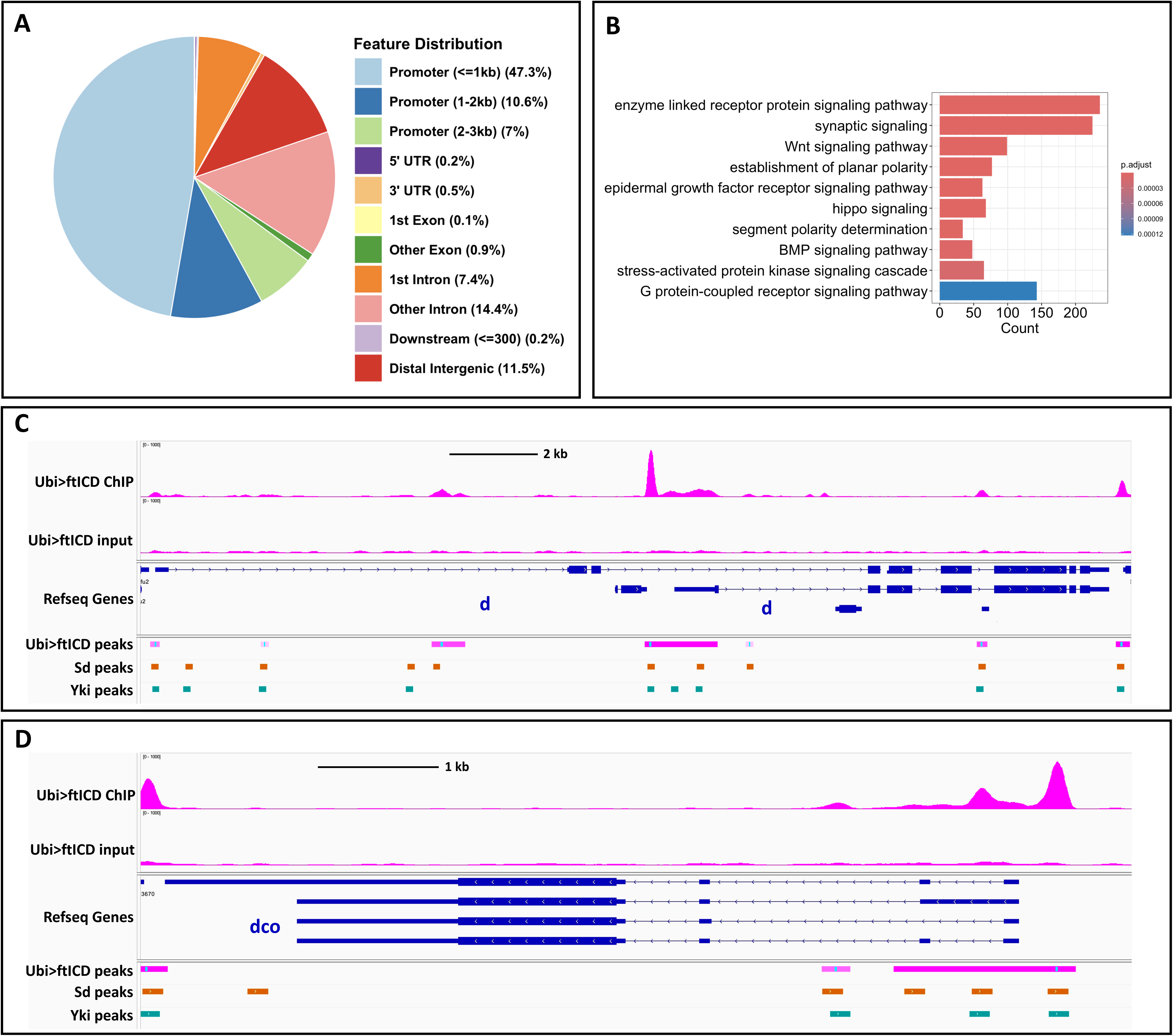
Genomic targets of overexpressed FtICD in larval eye-brain complexes. ChIP-seq was performed on 3^rd^ instar larval eye-brain complexes of larvae carrying ubi-Gal4 and UAS-ftICD-3xFLAG transgenes. (A), Feature distribution of FtICD binding regions, showing a majority of peaks localizing to promoter regions. (B). Top ten GO term signaling pathway enrichment hits of eye-brain FtICD ChIP seq data. (C) and (D), Genome browser tracks of examples of putative Ft target genes, *dachs* (*d*) (C) and *discs overgrown* (*dco*) (D). Sd and Yki ChIP peaks from public datasets are shown for reference.

### Motifs of diverse DNA-associated factors are enriched in Ft ChIP-seq

Since Ft has no identified DNA binding motif, it likely requires protein-protein interactions to bind to DNA targets. Some of those partners should be transcription factors whose DNA binding motifs might be enriched in the Ft ChIP peaks. To identify such factors, we performed *de novo* motif discovery and known motif enrichment analyses (Fig. S5F). We were able to identify several enriched motifs that closely resemble the motifs of known DNA associated factors, including DNA replication-related element factor (DREF) and GAGA Factor (GAF, the product of the Trithorax-like gene (Farkas et al., 1994). DREF was originally identified as a transcription factor that coordinately regulates the expression of DNA replication and proliferation-related genes in *Drosophila* (Tue et al., 2017). Among the genes it has been found to regulate are two core components of the Hippo pathway, *hpo* and *wts* (Fujiwara et al., 2011; Vo et al., 2014). These genes are also putative target genes of Ft ChIP-seq, and comparing the Ft and DREF ChIP peaks (Gurudatta et al., 2013) on *hpo* and *wts* revealed that the same regions were bound by the two factors, suggesting the potential co-regulation of *hpo* and *wts* by Ft and DREF (Fig. S5G).

GAF is a multifunctional protein long studied for its role in gene activation (Biggin & Tjian, 1988), but there is also evidence supporting its involvement in Polycomb-dependent repression (Busturia et al., 2001; Horard et al., 2000; Poux et al., 2001). GAF also acts as a barrier maintaining nucleosome-free regions (Adkins et al., 2006). A previous study showed that the transcriptional co-repressor Atrophin (Atro), which has been identified as a protein interactor of Ft (Fanto et al., 2003), is a major cofactor of GAF (Yeung et al., 2017).

Re-analysis of publicly available datasets of ChIP seq in eye discs (Ikmi et al., 2014; Oh et al., 2013) also revealed that Yki and Sd ChIP-seq (the Hippo pathway transcription cofactor and transcription factor, respectively) were highly correlated with Ft ChIP-seq. This suggested that a large fraction of genomic binding sites of nuclear Ft might be shared with Yki and/or Sd. An example for such a locus is the Hippo pathway component *expanded* (*ex*), where FtICD, Sd and Yki ChIP peaks perfectly align with each other (Fig.6A). In addition, Ft ChIP-seq was highly correlated with unphosphorylated RNA Pol II (Ikmi et al., 2014; Phatnani & Greenleaf, 2006), the active-enhancer marker H3K27Ac (Creyghton et al., 2010; Loubiere et al., 2016) (not shown) and ATAC-seq (Davie et al., 2015) (Fig.6B). In all these cases, the ChIP/ATAC-seq peaks can be grouped into subsets that are unique to FtICD (top section in Fig. 6B), common between FtICD and the other factors (middle section), or unique to the other factors (bottom section). Interestingly, the signal intensity tends to be highest in the common peaks. Overall, these results indicate that FtICD, Yki and Sd bind to many of the same genomic loci, which are also bound by RNA polII and correspond to open chromatin, suggesting that these loci are co-ordinately transcriptionally regulated by Ft, Sd and Yki.

**Fig. 6:**
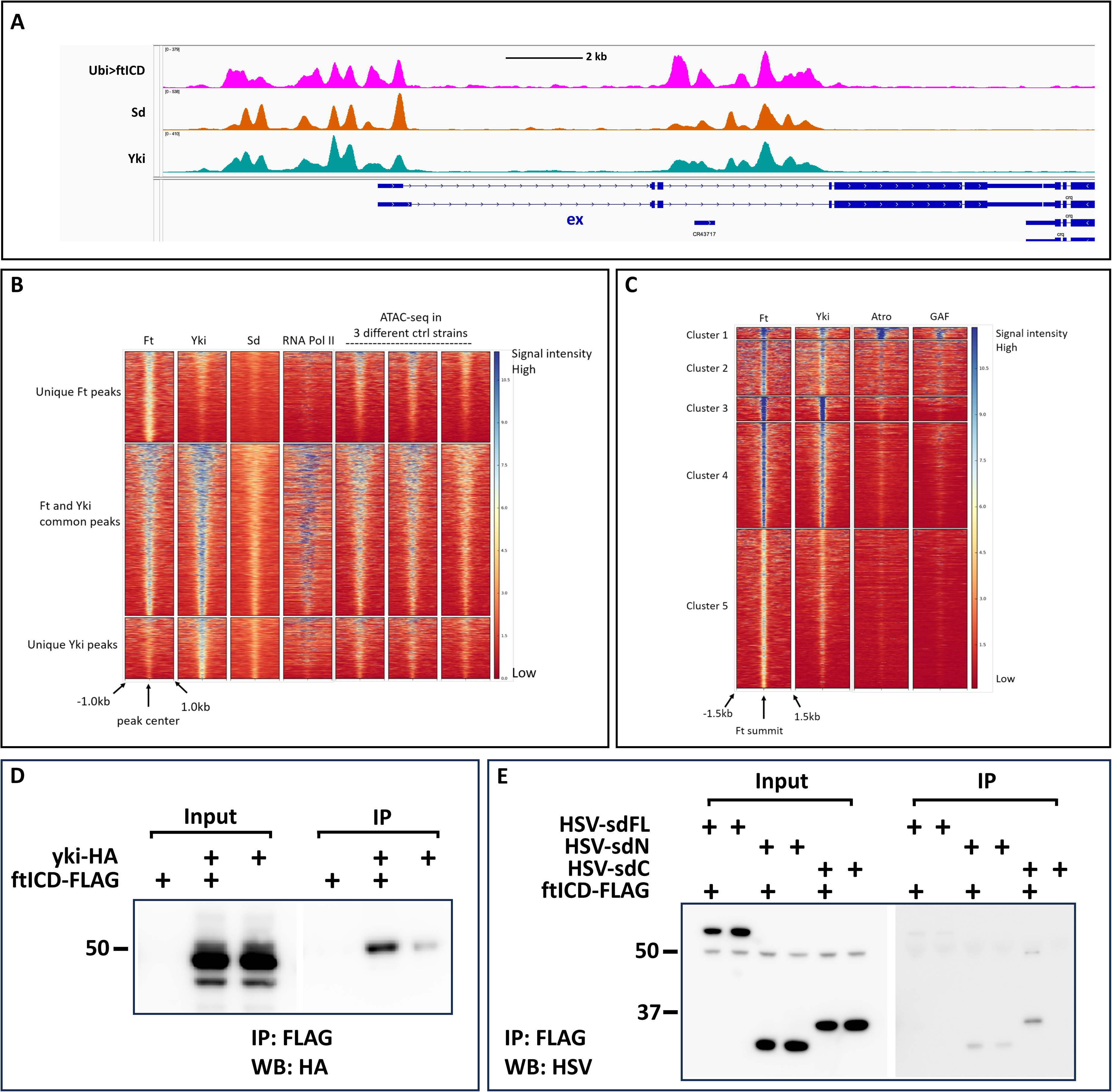
FtICD shares putative targets and physically interacts with Scalloped and Yorkie. (A) Genome browser tracks from eye-brain disc ChIP seq experiments showing the *expanded* (*ex*) locus. Ft peaks overlap with both Sd and Yki peaks. (B) Heatmaps comparing Ft, Yki, Sd and RNA Pol II ChIP seq, and ATAC peaks. (C), Heat maps comparing Ft, Yki, Atro and GAF ChIP seq peaks. (D) and (E) Co-immunoprecipitation experiments with FtICD. The indicated tagged proteins were overexpressed in S2 cells and lysates subjected to immunoprecipitation with anti-FLAG beads. The presence of co-immunoprecipitated proteins was analyzed by western blotting with anti-HA (D) or anti-HSV (E) antibodies. FtICD binds both Yki (D) and the Sd C-terminus (E).

When overlapping the ChIP seq datasets of FtICD, Yki and GAF, along with its partner Atro, we were able to subdivide the peaks into several clusters based on their similarity. One of the clusters comprises a subset of strong FtICD/Yki peaks that correlates with both strong Atro and GAF peaks (cluster 1 in Fig.6C), suggesting that there is a set of genes regulated co-ordinately by Ft, GAF and Atro.

The observation that FtICD, Sd and Yki bind to many of the same genomic loci prompted us to investigate whether these factors physically interact. To this end, we performed co-immunoprecipitation experiments using transiently transfected S2 cells. We found that FtICD binds specifically to Yki (Fig.6D) as well as the C-terminal part of Sd (Fig.6E). The C-terminal part of Sd has also been shown to bind Yki (Goulev et al., 2008a). Presently, it is not clear whether Sd can interact with both FtICD and Yki at the same time in a protein complex, or whether FtICD and Yki are alternative Sd binding partners. Additional layers of control, such as the presence or absence of other factors depending on developmental timing or tissue specificity, or the specific context of the binding site, could determine the identity of the proteins binding to these loci. However, these results suggest that Sd may be one of the transcription factors mediating Ft’s interaction with DNA.

### Potential targets of endogenous Ft protein

These data sets and analysis shown above point to a role for Ft in transcriptional regulation of target genes, including genes in pathways that involve Ft itself. However, these experiments were done under conditions of overexpressed Ft intracellular domain. In order to determine whether endogenous nuclear Ft can function in nuclear regulation, we made use of 2 fly strains carrying endogenously labeled Ft, named Ft-GFP^C^ and Ft-GFP^int^. These have, respectively, a C-terminally located GFP tag (Hale et al., 2015) or a novel tagged line we created with CRISPR in which GFP was inserted in the middle of the cytoplasmic domain (schematically represented in Fig.7A). Both strains are viable and fertile, indicating that these insertions do not interfere with Ft function.

**Fig. 7:**
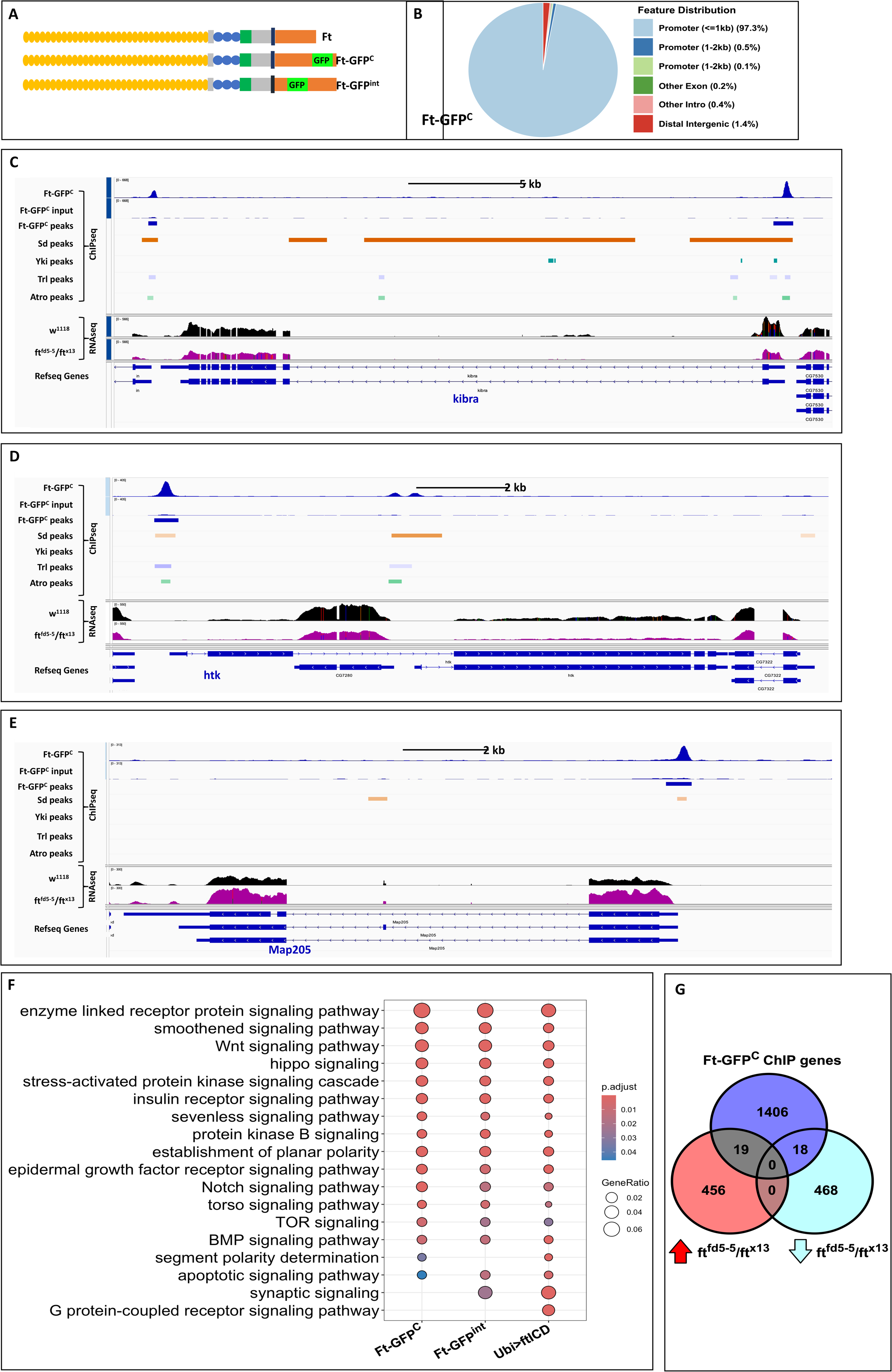
Genomic targets of endogenous ft protein in embryos. ChIP analysis was performed on 16-20hr old embryos with endogenously GFP-tagged ft, as well as on embryos overexpressing FLAG-tagged ftICD. (A) Schematic of CRISPR engineered strains used for endogenous ft ChIP experiments. Ft-GFP^C^ is a C-terminal fusion of GFP; GFP^int^ has the GFP sequence inserted in the HpN domain. (B) Feature distribution of Ft-GFP^C^ binding regions, showing a majority of peaks localizing to promoter regions. (C-E) Genome browser tracks of examples of putative Ft target genes, *kibra* (C), *hat-trick* (*htk*) (D), and Map205 (E). (F) GO analysis of signaling pathways among enriched biological processes. (G) Overlap of reproducible ChIP peaks in Ft-GFP^C^ embryos and differentially expressed genes in *ft* transheterozygous mutant embryos as determined by RNA seq. Red circle denotes genes upregulated in *ft* transheterozygotes, turquoise circle denotes downregulated genes.

Endogenously labeled Ft is readily detected in embryos in both strains (Fig. S7A), indistinguishable from endogenous Ft. We performed ChIP experiments on 16-20 hr old embryos from these strains, using a GFP antibody and again employing a 2-step crosslinking protocol. Peaks were selected using the same criteria as for the experiments using overexpressed FtICD. High-confidence peaks were defined based on reproducibility across three independent biological replicates. A total of 1,604 peaks were identified for Ft-GFP^C^ and 1,478 peaks for Ft-GFP^int^ (q-value < 0.01). The genomic distribution of Ft-GFP^C^ (Fig. 7B) showed strong enrichment near transcription start sites (≤1 kb from TSS, 97.3%) and a smaller fraction in distal intergenic regions (1.4%). This suggests that Ft may act at both proximal regulatory elements and *via* long-range chromatin interactions. Gene ontology analysis of genes associated with Ft-bound regions revealed significant enrichment in processes related to signaling pathways (e.g., Hippo and Wnt signaling) (Fig.7F), nucleosome/chromatin assembly, and nervous system development and function (Fig. S7G) (FDR < 0.05). Examples of putative Ft targets are shown in Fig. 7 C, D and E, including the Hippo pathway component *kibra* (Fig. 7C).

### Putative Ft target genes are dysregulated in *ft* mutants

To determine the biological relevance of Ft binding to genomic loci, we asked whether the putative Ft targets show a change in expression in the absence of Ft. To this end, we performed RNAseq experiments using *ft* mutant and *wild-type* embryos. Differential expression analysis identified 961 genes significantly dysregulated in *ft* mutants (FDR < 0.05 and log2-fold change > 0.50), with equal numbers being up-*vs* down-regulated (Fig.7G). Overlapping these genes with Ft-bound genes from Ft ChIP-seq data reveals 37 genes which show both reproducible Ft ChIP peaks and dysregulation in *ft* mutants. The examples in Fig. 7C-E include RNAseq tracks from *wild-type* (*w^1118^*, black traces) and *ft* transheterozygous mutant (*ft^fd5-5^/ft^x13^*, purple traces) embryos and show no significant change in RNA levels for *kibra* (Fig. 7C), significant downregulation for *hat-trick* (*htk*) (Fig. 7D), and significant upregulation for Map205 (Fig. 7E).

To identify putative DNA-binding cofactors or recruitment elements, we performed *de novo* motif discovery and known motif enrichment analyses using HOMER. For both Ft-GFP^C^ and Ft-GFP^int^, the DNA replication-related element factor (DREF) motif was most significantly enriched (P-value = 1e-348 in Ft-GFP^C^, 62.54% of peaks; P-value = 1e-395 in Ft-GFP^int^, 71.00%) (Fig. 8A). Common peaks in Ft-GFP^C^ and Ft-GFP^int^ showed significant enrichment for DREF (P-value = 1e-354, 72.78%), as well as BEAF-32B (P-value = 1e-84, 13.97%), pan (P-value = 1e-39, 21.60%), and the E-box motif (P-value=1e-15, 7.62%). Certain motifs were uniquely enriched in only one dataset: M1BP(Zf) in Ft-GFP^C^ (P-value = 1e-129, 33.33%) and the giant (gt) motif in Ft-GFP^int^ (P-value = 1e-20, 33.47%). BEAF-32 is an insulator protein in *Drosophila* that plays a role in genome organization (Yang et al., 2012). In addition, a previous study suggests that BEAF-32 is required for the activity of the Hippo pathway in terminal differentiation of neuronal subtypes (but not tissue growth) (Jukam et al., 2016). Thus, Ft may be involved in chromatin structure together with BEAF-32, in addition to a function in transcriptional regulation. E-box binding transcription factors (EBTFs) are proteins that bind to specific DNA sequences called E-boxes, which are regulatory elements in the promoter and enhancer regions of genes. Many EBTFs belong to the basic helix-loop-helix (bHLH) family, such as MYC and CLOCK-BMAL1, known to affect chromatin accessibility and transcriptional activation or repression depending on cellular context (Gordân et al., 2013).

**Fig. 8:**
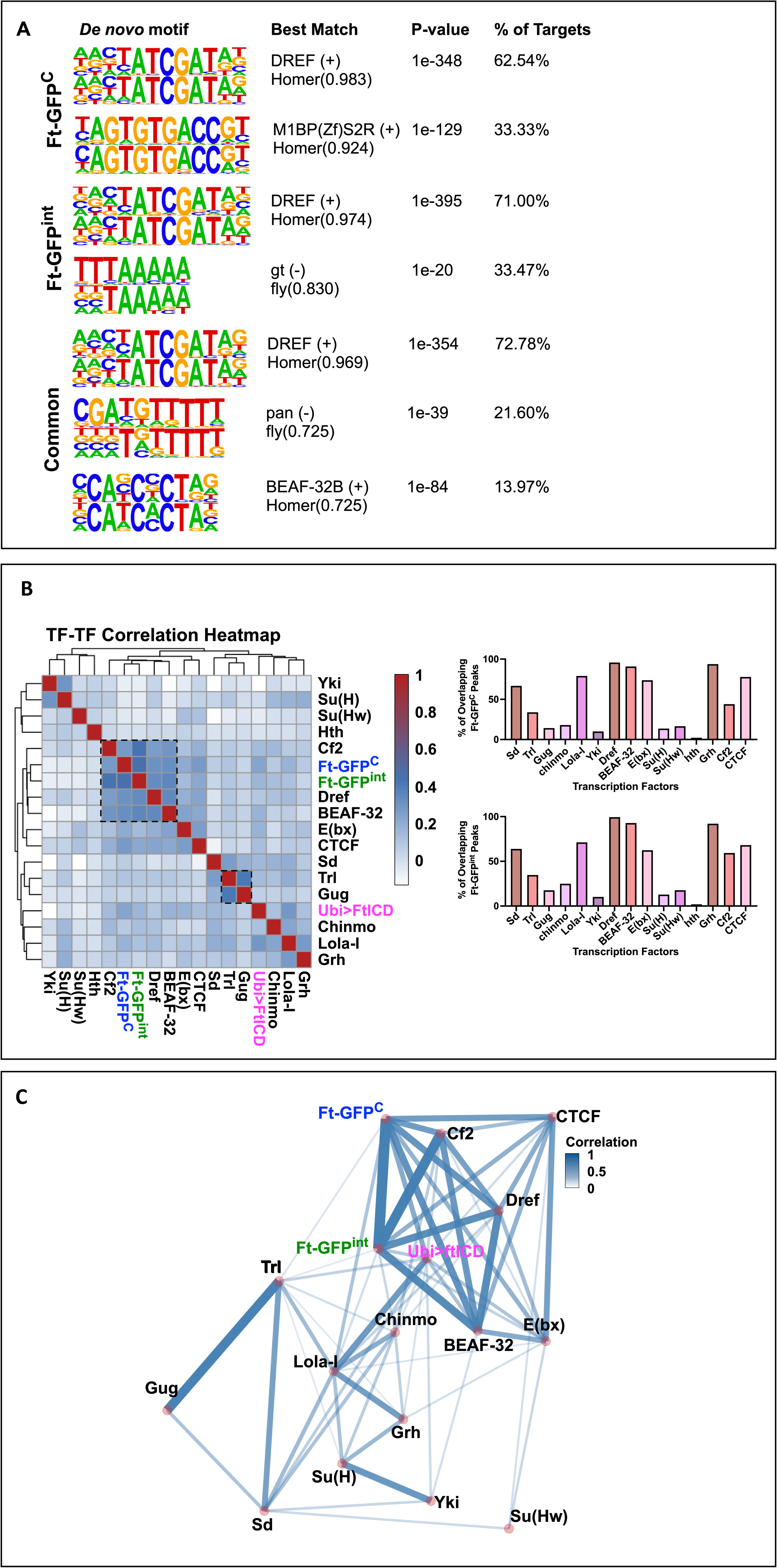
Identification of enriched motifs in Ft ChIP peaks. (A) Top results of *de novo* motif discovery in reproducible peak regions for Ft-GFP^C^, Ft-GFP*int*, and common peaks, sorted by the percentage of shared targets. (B) Correlation analysis of transcription factor binding in embryos using Ft-GFP^C^, Ft-GFP^int^ and a selected set of publicly available ChIP datasets. The bar graphs show the percentage of common peaks between Ft-GFP^C^ (top) and Ft-GFP^int^ (bottom) and the indicated transcription factors. (C) Proposed transcription factor network based on the correlation data in (B).

To explore potential co-regulatory interactions, we calculated pairwise Pearson correlation coefficients between Ft ChIP signal and publicly available ChIP-seq profiles from *Drosophila*. Datasets included developmental transcription factors (e.g., Sd, Yki, Lola-I, Grh), chromatin-associated proteins (e.g., CTCF, DREF, BEAF-32, E(bx), Trl), and other transcription factors (e.g., Gug, Chinmo, Su(H), Su(Hw), Hth, and Cf2). Correlation heatmaps revealed strong co-association between Ft-GFP^C^ or Ft-GFP^int^ peaks and factors such as DREF, BEAF-32, Lola-I, Grh, and CTCF (Nevil et al., 2020; Van Bortle et al., 2012), consistent with motif enrichment results. Network analysis using the igraph package (Fig. 8C) further revealed densely connected clusters centered on DREF, BEAF-32, E(bx), and CTCF, suggesting Ft participates in a regulatory module involving architectural proteins and transcriptional regulators.

Collectively, these data show that endogenously tagged Ft binds to thousands of genomic loci, preferentially near promoters, and that a subset of these loci show transcriptional dysregulation in *ft* mutants. The enrichment of motifs for known chromatin boundary and transcription factors –– particularly DREF and BEAF-32 –– together with co-binding correlation, suggests that Ft may regulate gene expression directly or through higher order of chromatin interactions during embryonic development.

## Discussion

The crucial roles of the Ft protein in several developmental pathways, namely Hippo and PCP, rely on its membrane localization, as demonstrated by the findings that a Ft protein lacking almost all of its extracellular portion but retaining the transmembrane domain is able to rescue *ft* mutants (Matakatsu & Blair, 2012), while the intracellular domain (ICD) alone is not. However, overexpression of the ICD does have mild effects on growth and patterning. We show here that FtICD can be readily detected in the nucleus when overexpressed, and we present evidence that it is able to bind specific genomic loci. We also show that FtICD is able to physically associate with the transcriptional co-activator Yki as well as with the DNA-binding transcription factor Sd, which together mediate downstream effects of the Hpo pathway. Although we have not been able to detect endogenous nuclear Ft protein by standard immunological methods, we demonstrate that Ft can be cleaved *in vivo* from a membrane-bound precursor and imported into the nucleus (Fig. 2). Furthermore, we show that endogenous Ft protein is able to bind to the promoter regions of specific genes, and that a subset of these putative Ft target genes are differentially regulated in *ft* mutants. As FtICD does not contain a known DNA-binding domain, we speculated that Ft acts as a transcriptional co-factor, requiring additional proteins to bind DNA and regulate the expression of its putative targets. The necessity of a 2-step crosslinking protocol for our ChIP experiments is consistent with this notion. By correlating our Ft ChIP-seq data with publicly available data for other transcription factors and chromatin regulators, we found strong correlations between Ft and several other factors such as DREF and BEAF-32, suggesting that Ft co-operates with these and other factors.

For our endogenous Ft experiments we made use of two different GFP-tagged Ft proteins (Ft-GFP^C^ and Ft-GFP^int^), which are both completely viable and fertile and indistinguishable from each other. However, although there is significant overlap between their binding targets, they also each show unique sites. This could indicate that the presence of GFP in different parts of the protein interferes differentially with FtICD’s binding to its different partners, which could in turn affect the specific loci to which the different complexes bind. Likewise, when we used overexpressed FtICD, we obtained a substantially larger number of peaks (almost 13,000 *vs* about 1,500 for endogenous Ft in embryos) which tended to be dispersed over the genome in a more non-specific manner, indicated by the peak distribution *vs* what is seen for endogenous Ft (Fig. 7 and Fig. S7), with a smaller proportion of peaks located at promoter regions. The fact that overexpressed FtICD is present in the cell at many-fold higher levels than endogenous Ft, as well as the absence of presumptive additional layers of regulation (i.e., regulation of Ft cleavage) may explain this observed higher binding.

We utilized previously published datasets for a select group of transcription and chromatin regulatory factors to explore our ChIP-seq results. Several factors were found to strongly correlate with Ft binding, including the transcription factor Sd and the co-activator Yki, an effector of the Hpo pathway. Interestingly, we found that both these proteins can bind to FtICD in S2 cells, raising the possibility that nuclear Ft functions in a protein complex that incudes these factors. We do not currently know whether all 3 proteins can interact to form a complex, or whether they are alternative binding partners. Sd has been shown to bind Yki *via* its C-terminal half (Goulev et al., 2008b), the same domain that we find to bind FtICD (Fig. 6). It is possible that one of the proteins acts as a bridge to mediate the interaction between the other two, or that binding of one precludes binding of the other. Future experiments will be needed to distinguish between these possibilities, but our observations suggest that one of the functions of nuclear Ft may be to modulate the activity of the Sd-Yki complex downstream of the Hpo pathway.

In our ChIP-seq experiments we find enrichment of a number of signaling pathways among the putative Ft targets. Significantly, this includes the Hpo as well as the PCP pathways. It has been shown that upstream Hpo pathway components such as *expanded* (*ex*), *kibra* and *Merlin* (*Mer*) are themselves targets of Hpo signaling (Genevet et al., 2010; Hamaratoglu et al., 2006), and our finding that FtICD binds to regulatory regions of these genes suggests the possibility that transcriptional regulation of these components by Ft may contribute to this feedback loop. Our finding that nuclear-localized FtICD can regulate gene transcription suggests that Ft can regulate the Hpo and PCP pathways at multiple levels: at the cell membrane *via* its well-characterized interaction with upstream components, and at the level of transcriptional control which may serve to fine-tune the activity levels of the pathways to ensure optimal control over crucial developmental processes.

## Materials and Methods

### Cloning of full-length ft-HA, ftICD-HA and ftμECD-HA

(from Seth Blair?) The UAS-ftC-3xFLAG construct was made by PCR amplifying ftC from pUASt-ftμECD, digesting with EcoRI and BglII and ligating to pUASt-attB. 3xFLAG was amplified from an Otefin-3xFLAG plasmid (Yonit Tsatskis, pers. comm.), digested with BamHI and BglII, and ligated to pUASt-attB-ftC. A stop codon was included in the primer used for amplification.

### Fly strains and genetics

UAS-ftFL-HA, UAS-ftICD-HA, UAS-ftμECD-HA UAS-ftC-3xFLAG was generated by site-specific insertion using the attP2 (chr3 68A) landing site (injections were performed by BestGene Inc). Transgenes were expressed using ubi-Gal4 (Bloomington, #32551) or GMR-Gal4 (). The HRE and HREm6 strains were a gift from D. Pan (S. Wu et al., 2008). To analyze HRE and HREm6 in *hpo* mutant clones they were crossed into FRT42D hpo^42-47^, a *hpo* null mutation (Wu et al., 2003, fly strain obtained from DJ Pan), or into FRT42D hpo^42-47^; UAS-ftC-3xFLAG. To generate the clones, these flies were crossed to the MARCM42D strain (hsflp UAS-mCD8-GFP; FRT42D tub-Gal80; tub-Gal4/TM6B) and larvae were heat shocked at 37°C for 30 min about 50 hrs after egg deposition.

CRISPR Ft-GFP^C^ and Ft-GFP^int^ Ft-GFP^C^ was a gift from David Strutt (Hale et al., 2015). For Ft-GFP^int^, the GFP moiety was inserted into the HpN domain.

### Tissue culture and co-immunoprecipitation

S2 cells were cultured at 25°C in S2 media with 10% fetal bovine serum, 100 u/ml penicillin and 100 μg/ml streptomycin. Cells were transfected using Effectene (Qiagen 301427) or Lipofectamine 3000 () according to the manufacturers’ protocols. For co-immunoprecipitation experiments, plasmids pAc-ftICD-3xFLAG, pAc-yki-HA (from??), pAc-HSV-sdFL, pAc-HSV-sdN, or pAc-HSV-sdC (gifts from DJ Pan) were transfected into 1.8×10^6^ cells per sample for 2 days. Cells were harvested, washed once in PBS and incubated in lysis buffer (100 mM KCl, 20 mM HEPES pH7.5, 5% glycerol, 2.5 mM EDTA, 0.2% IGEPAL CA-630, 1x protease inhibitor cocktail (Pierce, A32965), 1mM PMSF) for 5 min on ice. Lysates were cleared by centrifuging at 18,000xg for 15 min and incubated with anti-FLAG beads (Millipore, M8823) at 4°C overnight. Beads were washed 5×5 min at 4°C with lysis buffer containing 0.2% IGEPAL (for HSV-sd) or 0.4% IGEPAL (for yki-HA), and eluted for 5 min at 95°C in Laemmli buffer (BioRad, 1610737) containing 5% β-mercaptoethanol. Eluted proteins and inputs (5% of total) were analyzed by Western Blotting using anti-HA (Cell Sign Tech, C29F4, 1:2,000)) or anti-HSV (Invitrogen, PA5-116403, 1:2,000) followed by goat anti-rb (ProteinTech SA0001-2, 1:5,000) antibodies, and imaged using a BioRad ChemiDoc imager.

### Immunofluorescence

Larval tissues (from 3^rd^ instar wandering larvae) were dissected in ice-cold PBS and fixed in fresh 4% formaldehyde (FA, Thermo Scientific 28906) in PBS for 20 min with gentle agitation. Embryos were dechorionated in 50% bleach, rinsed, and fixed in 4% FA in PBS with heptane under vigorous shaking for 20 min. Vitelline membranes were removed by shaking with MeOH for 1 min, followed by re-hydration in PBST. After washing in PBST (PBS plus 0.1% Tween20) and blocking in PBST with 10% Normal Donkey Serum (Milllipore Sigma D9663), tissues were incubated with primary antibodies in blocking solution overnight at 4°C. After washing with PBST, tissues were incubated with secondary antibodies in blocking solution for 2 hrs at room temperature, washed in PBST, and in some cases incubated in Hoechst 33342 (Invitrogen H3570) 20μg/ml, for 5-10 minutes. Antibodies used were rat anti-ft (homemade) 1:500, mouse anti-HA (Roche, 12CA5), rat anti-HA (Roche 11867423001), mouse anti-RNA PolII Ser2P (H5, abcam ab24758), mouse anti-Lamin B (DSHB ADL67.10), rabbit anti-βgal (MP Biomedical 8559761) 1:500, rabbit anti-GFP (Chromotek pabg1 or Abcam ab290) 1:500, mouse anti-FLAG (M2, Sigma F1804) 1:500. Secondary antibodies were Alexa488, Alexa568, Alexa594 or Alexa647-coupled and obtained from Invitrogen.

### X-gal staining

Larval tissues (from 3^rd^ instar wandering larvae) were dissected and fixed in 1% glutaraldehyde for 5 min at room temperature, washed with PBST and then washed for 30 min with staining solution (10 mM Na phosphate buffer pH 7.2, 150 mM NaCl, 1 mM MgCl_2_, 3.1 mM K4[FeII(CN)6], 3.1 mM K3[FeII(CN)6], 0.3% Triton X-100). The staining was developed for 20 min with 2 mg/ml X-gal in staining solution, and washed with PBST.

### NLS and NES prediction

For NLS and NES prediction, the following online computational tools were used with default parameters: NLStradamus (revision r.9) (Nguyen Ba et al., 2009), cNLS Mapper (last update: 2012/11/7) (Kosugi et al., 2009), NucPred (Brameier et al., 2007), NetNES (version 1.1) (La Cour et al., 2004), NESsential (Fu et al., 2011). The amino acid sequence of FtICD (^4610^RFRGKQ-EEYV^5147^) was used as input sequence. The molecular weight of FtICD was calculated using “Protein Molecular Weight” in bioinformatics.org (Stothard, 2000).

### RNA-seq analysis

Genotypes used were w^1118^, ft^fd5-5^/Cyo[Dfd-GFP] and ft^x13^/CyO[Dfd-GFP]. Embryos were collected for 4 hrs, aged for 16 hrs at 25°C, and dechorionated in 50% bleach. 200 embryos per sample were collected, transferred to 250 μl of LBA buffer, flash frozen in lN_2_ and stored at –80°C. For *ft* transheterozygotes, GFP-negative embryos were selected. RNA was prepared using the ReliaPrep RNA Tissue Miniprep System (Promega, Z6111) according to the manufacturer’s instructions. RNA-seq libraries were prepared with 10 ng of total RNA using the SMARTer Ultra Low RNA kit for Illumina Sequencing (Takara-Clontech) per manufacturer’s protocol. cDNA was fragmented using a Covaris E220 sonicator using peak incident power 18, duty factor 20%, cycles per burst 50 for 120 seconds. cDNA was blunt ended, had an A base added to the 3’ ends, and then had Illumina sequencing adapters ligated to the ends. Ligated fragments were then amplified for 12-15 cycles using primers incorporating unique dual index tags. Fragments were sequenced on an Illumina NovaSeq X Plus using paired end reads extending 150 bases.

Quality of raw reads was assessed using FastQC (v.0.10). Adapter sequences and low quality or short reads were removed using Trimmomatic (v.0.33) (Bolger et al., 2014). Using Salmon v1.8.0 (Patro et al., 2017), high-quality reads were pseudo-aligned and quantified to the *D. melanogaster* reference transcriptome (dm6, BDGP6.28). Corresponding GTF annotation files (Drosophila_melanogaster.BDGP6.22.98.gtf) were obtained from Ensembl. Computational preprocessing was conducted using resources provided by Washington University School of Medicine in St. Louis.

Differential gene expression (DGE) was performed using the edgeR package (Robinson et al., 2009). Low-expressed genes were filtered using edgeR’s default independent filtering. Genes with an FDR < 0.05 and |log2 fold change| > 0.50 were defined as differentially expressed genes (DEGs).

Functional enrichment of DE genes was performed using clusterProfiler v.4.10.1 (T. Wu et al., 2021). Gene Ontology (GO) terms and pathways with Bonferroni-corrected *P*-values < 0.05 were considered significantly enriched.

### ChIP-seq analysis

Chromatin immunoprecipitation followed by sequencing (ChIP-seq) was performed to identify genomic regions bound by Ft-GFP^int^ and Ft-GFP^C^ fusion proteins in *Drosophila melanogaster* embryos and larval eye-brain complexes. The following genotypes were used: Ft-GFP^C^, Ft-GFP^int^, Ubi-Gal4; UAS-FtC-3xFLAG, and *w^1118^*. All procedures were performed according to ENCODE consortium guidelines for ChIP-seq experiments (Landt et al., 2012).

Embryos were collected for 4 hrs, aged for 16 hrs at 25°C, and dechorionated in 50% bleach. For 2-step crosslinking they were transferred to scintillation vials containing 7 ml of heptane and 2.5 ml buffer A (60 mM KCl, 15 mM NaCl, 4 mM MgCl_2_, 15 mM HEPES pH 7.5) with 2 mM EGS (Thermo Scientific 21565) and shaken at 300 rpm for 45 min. The aqueous phase was removed and replaced with 3 ml buffer A containing 1.8% formaldehyde (Thermo Scientific 28906), and shaken again at 300 rpm for 15 min. Crosslinking was stopped by adding 400 μl of 2.5 M glycine and gently rotating for 5 min. Embryos were washed several times with buffer A containing 0.1% Triton X-100, transferred to Eppendorf tubes in approximately 50 μl aliquots, flash frozen in liquid nitrogen and stored at –80°C.

Eye-brain complexes (100 per genotype) from wandering 3^rd^ instar larvae were dissected in ice-cold PBS and crosslinked in 1 ml PBS containing 2 mM EGS for 45 min at room temperature with gentle agitation, washed with PBS, followed by crosslinking with 1.8% formaldehyde in PBS for 15 min and quenching with 250 mM glycine for 5 min. Tissues were flash frozen in liquid nitrogen and stored at –80°C.

Material was resuspended in 1 ml ice-cold lysis buffer 1 (140 mM NaCl, 15 mM HEPES pH 7.5, 1 mM EDTA, 0.5 mM EGTA, 1% Triton X-100, with freshly added 0.5 mM DTT, 0.1% Na-deoxycholate, and 1x protease inhibitor cocktail (Pierce A32965)) and homogenized with 15 strokes in a 2ml glass Dounce homogenizer (Kontes 885303-0002). Nuclei were pelleted by centrifugation for 5 min at 4000g, 4°C, and lysed in 600 μl nuclear lysis buffer (lysis buffer 1 plus 0.1% SDS and 0.5% N-lauroylsarcosine) for 30 min at 4°C with rotation. DNA was sheared by sonication using a Bioruptor (Diagenode) at High Intensity setting, for a total of 10-12 30 sec on – 30 sec off pulses at 4°C, followed by centrifuging for 5 min, 14,000g at 4°C. 1% of the resulting supernatant was removed as input sample and kept at –20°C. Anti-GFP (rabbit, abcam 290, 1 μl) or anti-FLAG (mouse, M2, Sigma F1804, 5 μg) antibodies were added to the chromatin and incubated with gentle agitation overnight at 4°C. Equilibrated MagnaChIP A+G beads (EMD Milllipore 16-663, 30μl per sample) were added for 4 hrs at 4°C, followed by 10 min washes in low salt buffer (150 mM NaCl, 20 mM Tris pH 8.0, 2 mM EDTA, 1% Triton X-100, 0.1% SDS, 1x protease inhibitor), high salt buffer (same as low salt buffer, except 500 mM NaCl), LiCl buffer (250 mM LiCl, 10 mM Tris pH 8.0, 1 mM EDTA, 1% IGEPAL CA630, 1% Na-deoxycholate, 1x protease inhibitor), and TE (10 mM Tris pH 8.0, 1 mM EDTA, 1x protease inhibitor), all at 4°C with rotation. ChIP and input samples were de-crosslinked and eluted from the beads by incubation in elution buffer (50 mM Tris pH 8.0, 10 mM EDTA, 1% SDS, 200 mM NaCl) at 65°C with 900 rpm shaking in a thermomixer (Eppendorf) overnight. Eluted DNA was purified by RNaseA and proteinase K digestion followed by phenol/chloroform extraction and EtOH precipitation. Library preparation and sequencing (standard paired-end reads 2×150 on NovaSeq6000 or NovaSeq S4) was performed by GTAC (https://gtac.wustl.edu/).

Raw sequencing reads were processed using Trimmomatic v0.33 (Bolger et al., 2014) to remove low-quality bases and adapter contamination, employing standard trimming parameters. Alignment of trimmed reads to *Drosophila melanogaster* reference genome (dm6, BDGP6.28) was performed by BWA v0.7.16a tool (Li & Durbin, 2009). BAM files were created by SAMtools 1.6 (Li et al., 2009), and genome coverage (BigWig) files were generated by makeUCSCfile.pl (HOMER v4.11.1) (Heinz et al., 2010).

Peak calling was performed using MACS2 v2.1.2 for paired-end ChIP-seq data (Y. Zhang et al., 2008). For each genotype, biological replicates were processed individually before reproducibility assessment.

To retain reproducible peaks (high-confidence peaks), multiIntersectBed (bedtools v2.30.0) (Quinlan & Hall, 2010) was used. For Ft-GFP^C^ and Ft-GFP^int^, peaks present in at least three independent replicates were retained as reproducible binding regions. To further eliminate background signal, we used the *w^1118^* input as a negative control. Peaks overlapping with *w^1118^* were excluded unless they had a log₂ fold enrichment > 1, in which case they were retained as potential true positives. The final set of high-confidence reproducible peaks comprised both (1) peaks unique to Ft-GFP^int^ or Ft-GFP^C^, and (2) overlapping peaks meeting the fold change threshold.

For peak annotation, ChIPseeker v1.38.0 (Y. Zhang et al., 2008) with the annotatePeak function was used to associate reproducible peaks with genomic features. Feature distribution across promoter, intronic, exonic, and intergenic regions was also extracted using ChIPseeker.

Gene Ontology (GO) and KEGG pathway enrichment analyses were conducted on genes associated with reproducible peaks using clusterProfiler v4.10.1 (T. Wu et al., 2021). Only terms with a Bonferroni-adjusted *P* value < 0.05 were considered significantly enriched.

Motif analysis was conducted using HOMER default settings to discover enriched motifs within reproducible peak regions for Ft-GFP^C^ and Ft-GFP^int^. Only motifs associated with peaks having a score > 1000 were retained for downstream analysis.

For visualization, feature distribution plots were generated using ggplot2 v3.5.1. Venn diagrams showing overlap of reproducible ChIP regions (between Ft-GFP^int^ and Ft-GFP^C^) and their intersection with differentially expressed genes from *ft* transheterozygous mutants were plotted using ggVennDiagram v1.5.2.

To examine potential coregulation and functional relationships among transcription factors (TFs), a pairwise TF–TF correlation analysis using publicly available ChIP-seq datasets (https://chip-atlas.org/) from *Drosophila* embryos was conducted. The analysis included TFs and chromatin regulators with defined embryonic ChIP-seq profiles: Sd, Trl, Gug, Chinmo, Lola-I, Yki, BEAF-32, E(bx), Su(H), Su(Hw), Hth, Grh, Cf2, and CTCF. In cases where embryo-specific data were unavailable (e.g., Dref), datasets from all available tissues and cell types were used.

All public ChIP-seq datasets included were filtered to retain high-confidence binding peaks with Q-values < 1 × 10⁻⁵. Peak overlaps between TFs were quantified, and pairwise Pearson correlation coefficients were computed across binding profiles to assess TF co-binding relationships.

A heatmap of the resulting correlation matrix was generated using pheatmap v1.10.2, and transcriptional regulatory networks were visualized using igraph v2.0.3, facilitating interpretation of TF clustering and functional associations.

### Wing analysis

Strains used were enGal4; UAS-ftC-3xFLAG, wts^X1^/TM6B, and yki^B5^/CyO. To obtain heterozygous animals, these were crossed to w^1118^. All animals were grown at low density, and adults of the desired genotypes selected. Wings were removed, mounted in Euparal (Hindolveston) or Entellan new (Sigma, 1.07961.0500), and photographed on a Nikon SMZ1270 microscope equipped with a Nikon DS-F*i*3 camera, using Zen software. Images were analyzed and measurements obtained using ImageJ software (NIH).

## Acknowledgments

We thank D. Strutt and D. Pan for fly stocks. We are grateful to members of the McNeill lab for helpful discussion, and Alex Earl for technical assistance. Stocks obtained from the Bloomington Drosophila Stock Center (National Institutes of Health [NIH] P40OD018537) were used in this study. This work was funded by NIH R01GM138853.

