## supplemental figures for "Fat cadherin cleavage releases a transcriptionally active nuclear fragment to regulate target gene expression"

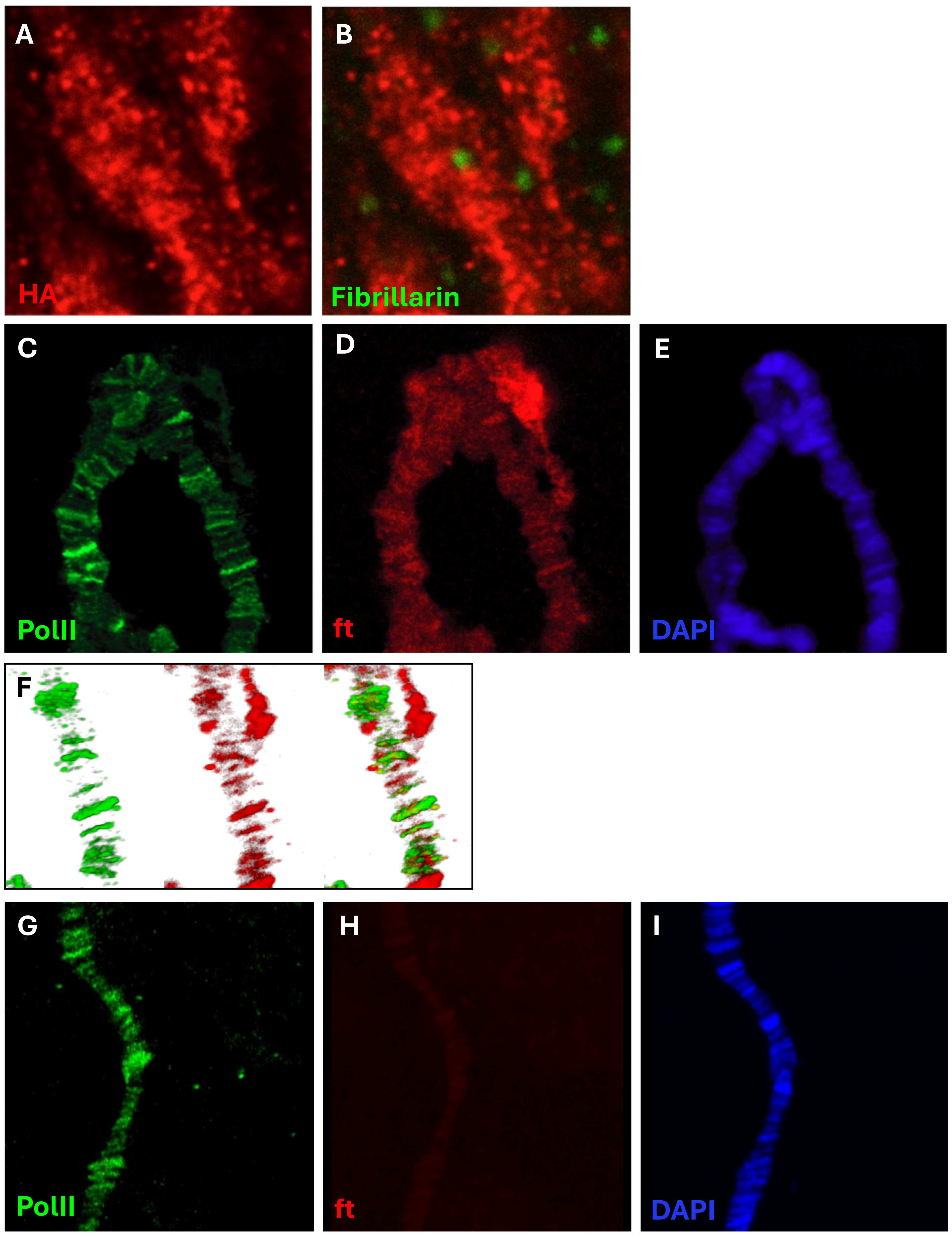


**Fig. S1:** (A-B) Wandering L3 larval eye disc expressing FtICD-HA under GMR-Gal4 control, stained for HA (red) and the nucleolus marker Fibrillarin (green). FtICD accumulates in the nucleus but is excluded from the nucleolus. (C-I) Salivary gland nuclei stained with antibodies to polII (green), ft (red) and with DAPI. (C-E), ft is seen localizing to chromosome bands in wild-type, but not in a ft mutant ($ft^{x13}$), (G-I).

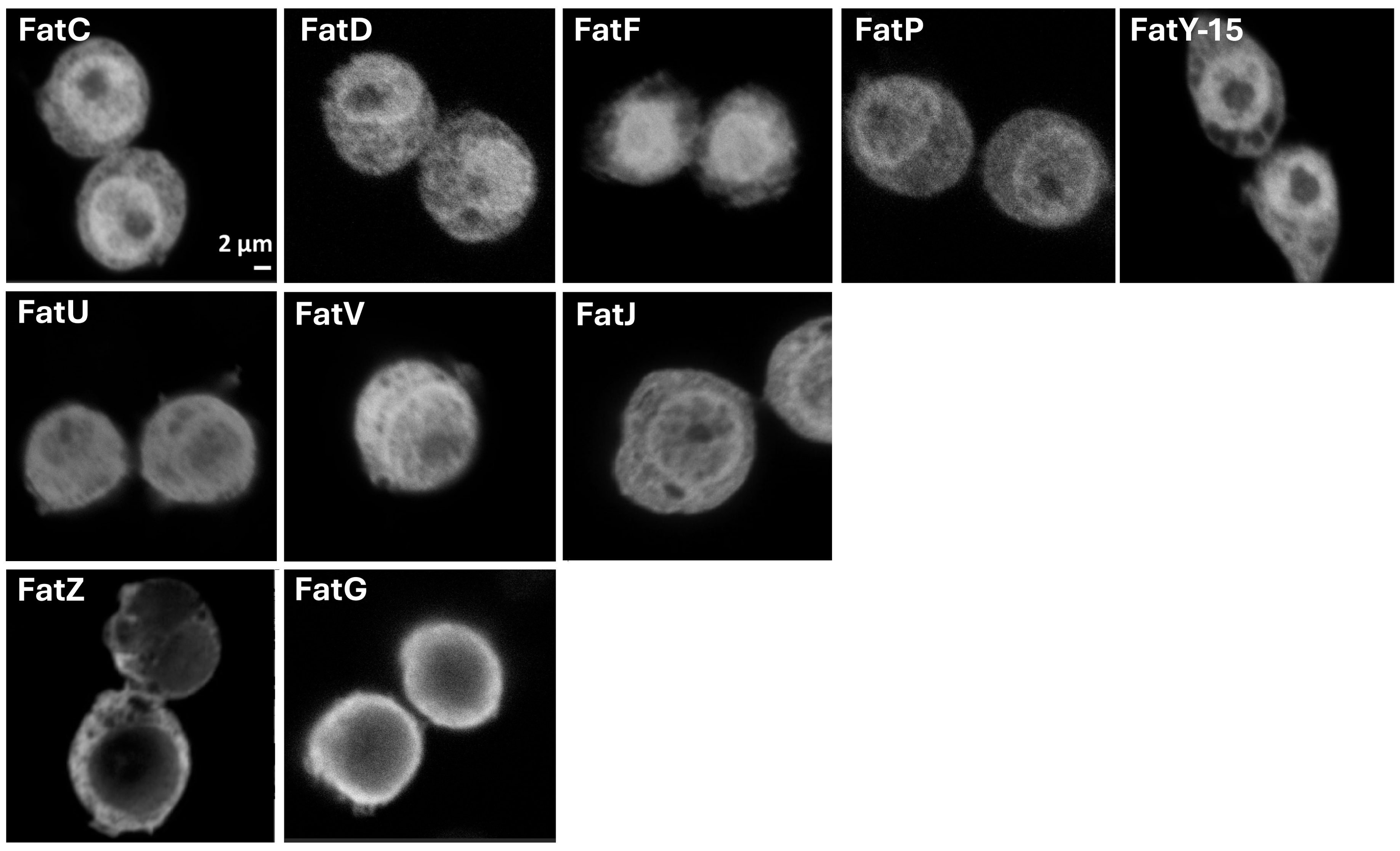


**Fig. S2: Identification of nuclear import and export sequences in ftICD**. Diagrams of the constructs used to transiently transfect S2 cells are in Fig. 2F. Images show cells immunostained with HA and are grouped by nuclear enriched localization (top row), ubiquitous localization (middle row), and nuclear excluded localization (bottom row).

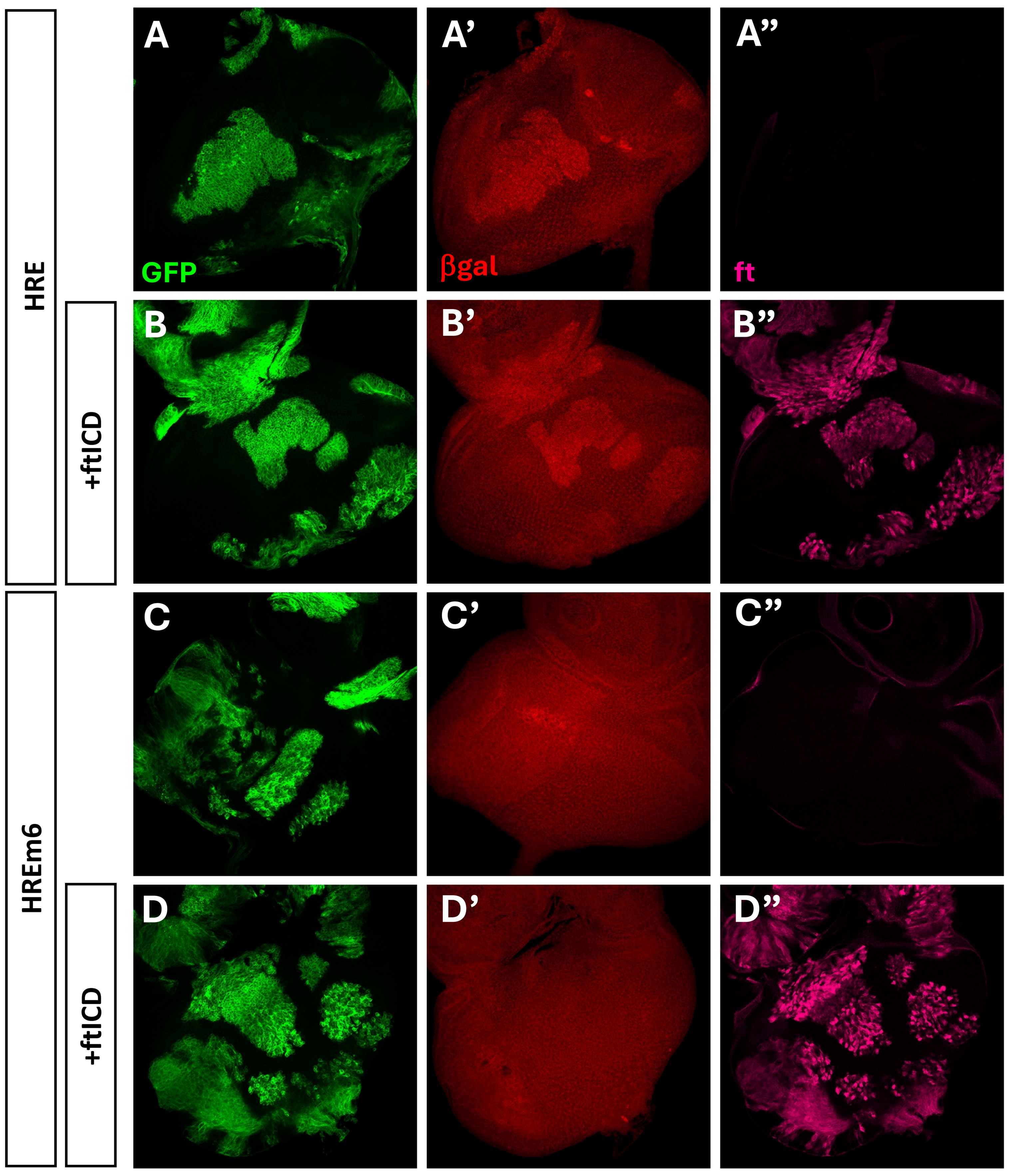


**Fig. S3: ftICD can not activate the hippo pathway target *diap1* in eye discs.** *hpo*$^{42\text{-}47}$ mutant clones (marked by GFP expression in green) were generated in eye discs in larvae carrying either the HRE (A, B) or HREm6 (C, D) reporter. Larvae also carrying the UAS-ftICD transgene are shown in (B) and (D). Discs were stained with antibodies to βgal (red) and ft (magenta). Wild-type HRE is activated in *hpo* clones (A, B), whereas HREm6, which lacks the sd binding site, is not activated in the absence (C) or presence of overexpressed ftICD (D).

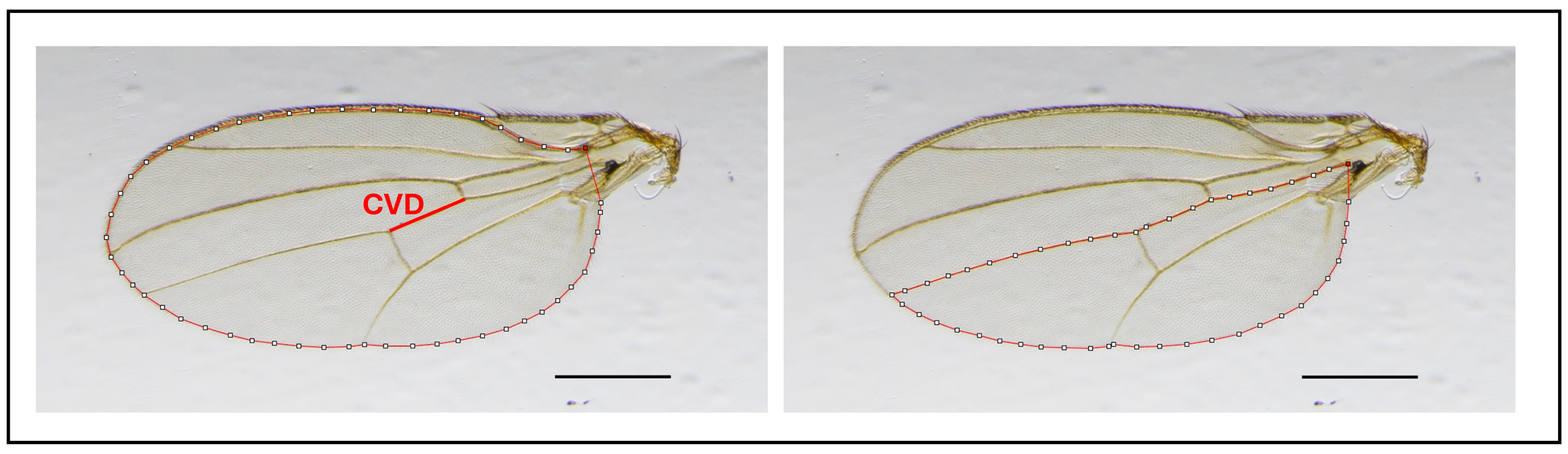


**Fig. S4: Wing parameter definitions.** (A) Total wing area and crossvein distance (CVD). (B) Posterior wing area, defined as area below L4. Scale bar, 500 μm.

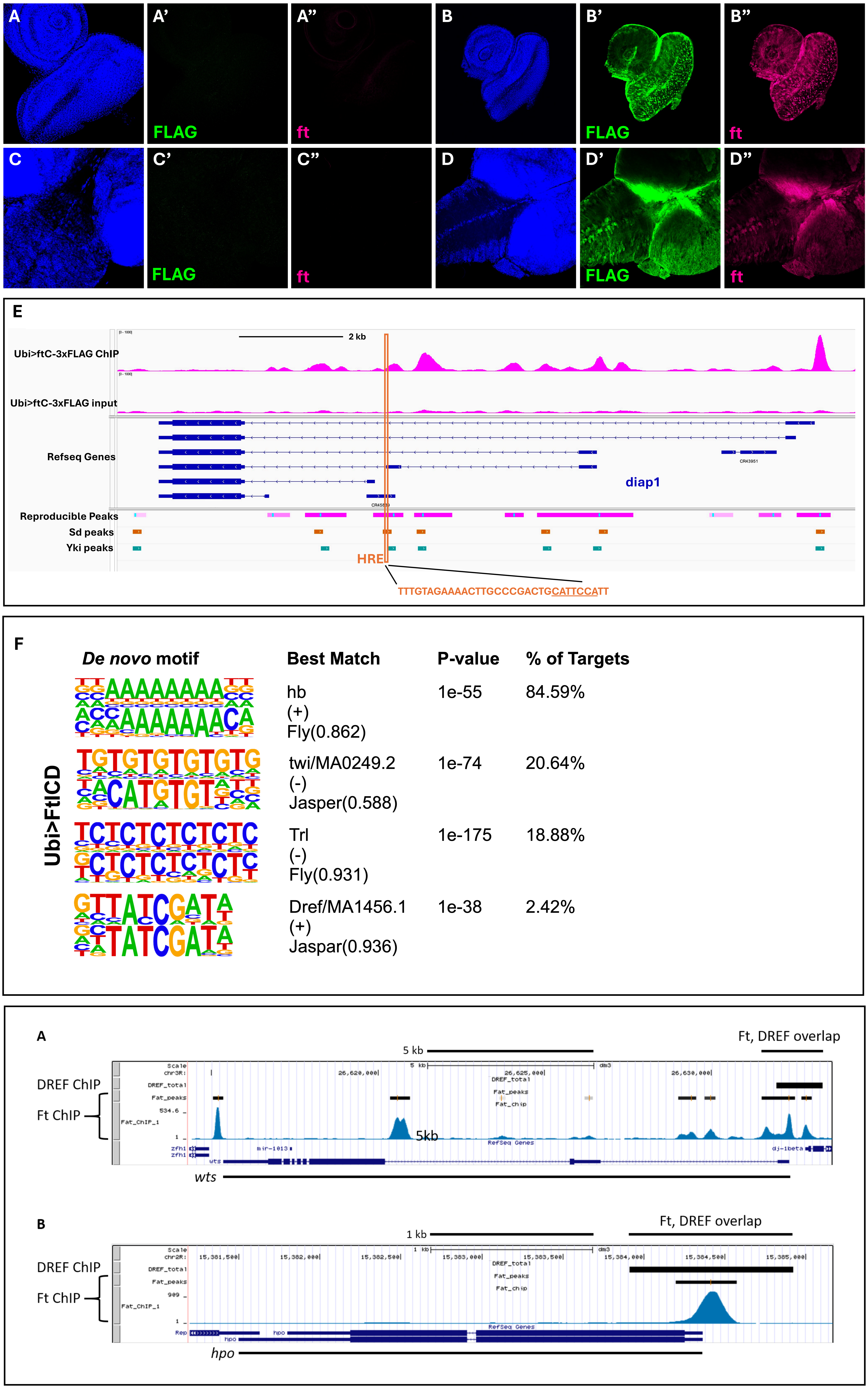


**Fig. S5: Genomic targets of Ft protein in larval eye-brain complexes.** (A-D), Expression of Ft in *wild-type* larvae (w[1118]) (A and C), and larvae carrying ubi-Gal4; UAS-ftC-3xFLAG (C and D) transgenes, stained with FLAG and Ft antibodies. (A) and (B) show eye discs, (C) and (D) show brains. Levels of Ft expression are strongly increased in the transgenic animals, and nuclear localization of ftICD can be seen. All images were taken with identical settings so the endogenous Ft expression in w[1118] is not visible with this exposure. (E), Genome browser track of the *diap1* locus. The orange box marks the location of the HRE (with the Sd binding site underlined) and shows that a ft ChIP peak overlaps this region. Sd and Yki ChIP peaks from public datasets are also shown for reference. (F) *De Novo* motif enrichment for eye-brain complex ChIP-seq of ubi>FtICD animals. (G) Genome browser tracks of FtICD and DREF ChIP peaks (from Gurudatta et al., 2013) on the *wts* and *hpo* loci.

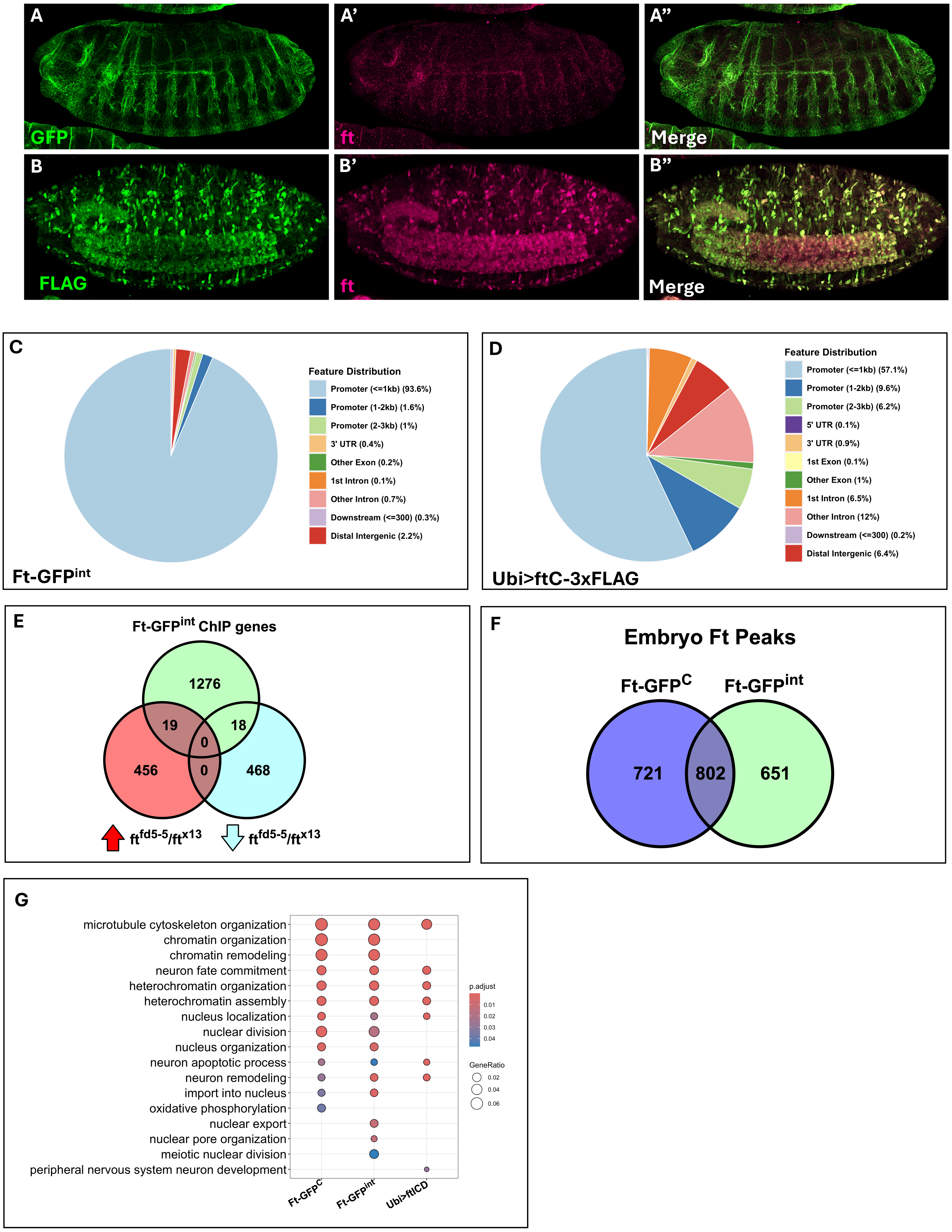


**Fig. S7: Endogenous tagged Ft protein and its genomic targets.** (A) GFP and Ft antibody staining of late stage embryos from the Ft-GFP$^{C}$ strain. The GFP expression faithfully represents endogenous Ft staining. (B) FLAG and ft antibody staining of embryos overexpressing ftICD-3xFLAG under ubi-Gal4 control. (C) Feature distribution of Ft-GFP$^{int}$. (D) Feature distribution of embryos overexpressing ftICD under ubiquitin-Gal4 control. (E) Overlap of reproducible ChIP peaks in Ft-GFP$^{C}$ embryos and differentially expressed genes in *ft* transheterozygous mutant embryos as determined by RNA seq. (F) Overlap between reproducible ChIP peaks in Ft-GFP$^{C}$ *vs* Ft-GFP$^{int}$ embryos. (G) GO term analysis for embryo Ft ChIP-seq focused on epigenetics related processes, neuronal development and nuclear structure and function.
